# Harnessing secretory pathway differences between HEK293 and CHO to rescue production of difficult to express proteins

**DOI:** 10.1101/2021.08.16.455786

**Authors:** Magdalena Malm, Chih-Chung Kuo, Mona Moradi Barzadd, Aman Mebrahtu, Num Wistbacka, Ronia Razavi, Anna-Luisa Volk, Magnus Lundqvist, David Kotol, Fredrik Edfors, Torbjörn Gräslund, Veronique Chotteau, Ray Field, Paul G. Varley, Robert G. Roth, Nathan E. Lewis, Diane Hatton, Johan Rockberg

**Author notes:** Corresponding authors: Johan Rockberg, KTH - Royal Institute of Technology, School of Engineering Sciences in Chemistry, Biotechnology and Health, S- 106 91 Stockholm, Sweden, Or: Nathan E. Lewis, University of California; San Diego, Departments of Pediatrics and Bioengineering; La Jolla; CA 92093; USA. These authors contributed equally to this work (Magdalena Malm, Chih-Chung Kuo).

## Abstract

Biologics represent the fastest growing group of therapeutics, but many advanced recombinant protein moieties remain difficult to produce. Here, we identify bottlenecks limiting expression of recombinant human proteins through a systems biology analysis of the transcriptomes of CHO and HEK293 during recombinant overexpression. Surprisingly, one third of the challenging human proteins displayed improved secretion upon host cell swapping from CHO to HEK293. While most components of the secretory machinery showed comparable expression levels in both expression hosts, genes with significant expression variation were identified. Among these, ATF4, SRP9, JUN, PDIA3 and HSPA8 were validated as productivity boosters in CHO. Further, more heavily glycosylated products benefitted more from the elevated activities of the N- and O-glycosyltransferases found in HEK293. Collectively, our results demonstrate the utilization of HEK293 for expression rescue of human proteins and suggest a methodology for identification of secretory pathway components improving recombinant protein yield in HEK293 and CHO.

## Introduction

The Chinese hamster ovary (CHO) cell line is commonly used for producing recombinant proteins (r-proteins) as it enables efficient expression of proteins with the need for human-like post-translational modifications. The CHO cell line provides several attractive properties for large-scale production of biopharmaceuticals, such as the ability to be cultivated at high cell densities in serum-free and chemically defined media and low risk of infection of human viruses (Kim et al., 2012). Currently, CHO cell lines are the biopharmaceutical mammalian workhorses, producing 84% of recently approved monoclonal antibodies (Walsh, 2018). However, with the boom of biologics within the pharma industry combined with more complex pharmaceutical proteins reaching the market, there is a demand for bioproduction platforms that can produce more difficult to express proteins. Data from the ongoing human secretome project (Tegel et al., 2020; Uhlén et al., 2019), a comprehensive research study that includes the secreted production of >1500 human proteins, suggests approximately 35% of proteins are challenging to efficiently produce in secreted form by the CHO expression system. For such difficult to express proteins, alternative expression systems based on other expression hosts or engineered cell lines may provide improved expression titers. As cells of different origins can have tissue-specific expression patterns of secretory pathway genes (Feizi et al., 2017), such differences can bring about variation in expression and processing of r-proteins depending on expression host, which in turn can impact the protein’s stability, function, activity, immunogenicity and production titer. Moreover, poor expression of human proteins in CHO has previously been overcome by exogenous expression of human endoplasmic reticulum (ER)-associated proteins in the CHO production cells (Le Fourn et al., 2014; Hansen et al., 2017). Thus one could speculate, when focusing on improving expression of many challenging human proteins, that the CHO cell line may have a secretory systems disadvantage compared to human production hosts. In particular, the human embryonic kidney 293 (HEK293) cell line has traditionally been a popular alternative for bioproduction and is commonly used for transient production of proteins for research and preclinical studies. HEK293 is considered easy to transfect, it adapts well to suspension cultivation, grows rapidly in serum-free media and is capable of producing r-proteins at high titers (Jiang and Zhu, 2013; Pham et al., 2006). In particular, glycosylation profiles of r-proteins produced in either CHO or HEK293 have been shown to differ (Böhm et al., 2015; Croset et al., 2012; Goh and Ng, 2018). In the context of biopharmaceutical production, specific demands for human post-translational modifications for certain r-proteins have made HEK293 successful alternatives to the conventional CHO cells and in 2018 five approved protein therapeutics were produced in HEK293 cell lines (Dumont et al., 2016; Walsh, 2018).

Here, we evaluated alternative expression systems for production of a set of human difficult to express secreted proteins or extracellular domains of single-pass plasma membrane-anchored proteins (Tegel et al., 2020; Uhlén et al., 2019). Expression systems were evaluated both based on r-protein expression levels but also systemic differences based on transcription of secretory pathway components between cell lines. Initially, a comparison of two CHO-based expression systems was carried out followed by a comparison of the secreted expression between CHO and HEK293. Moreover, transcriptome data from CHO and HEK293 cells transiently expressing proteins was analyzed to map differences in secretory pathway components between HEK293 and CHO. Based on the most profound differences in the expression of secretory pathway components between the cell lines, secretory pathway genes with significant impact on secreted productivity of human difficult-to-express proteins were identified. These include genes that assist protein secretion in a product-independent fashion. Additionally, we coupled specific post-translational modifications (PTMs) of different r-proteins with the protein titer improvements from CHO to HEK293 cells and showed that differences in titer improvements can be jointly explained by PTM features of the r-proteins and the activities of the enzymes responsible for these PTMs. These highly product-specific genes enable bespoke cell line designs that cater to the unique secretory requirements of different r-proteins, and allows for a more rational selection of cell hosts for a given r-protein.

## Results

### The two CHO platforms ExpiCHO and QMCF provide different benefits for difficult to express proteins

Due to differences in expression platform protocols, performance in productivity of r-proteins may vary even between hosts of the same origin. To shine light on differences between platforms for a range of difficult to express proteins, we compared expression levels in various systems. Initially, we evaluated two CHO-based expression systems by expressing a set of 23 challenging human proteins in the ExpiCHO system and compared secreted productivities to Human Secretome Project production data, wherein expression was performed using the episomal stable QMCF technology (Silla et al., 2005). Briefly, this system utilizes CHOEBNALT85 cells stably expressing the Epstein-Barr virus EBNA-1 protein and the mouse polyomavirus (Py) large T antigen, which facilitates nuclear retention and replication of the pQMCF expression vector. The ExpiCHO platform is a fully transient system that enables protein production at very high cell densities. In this comparison, transient expression of the 23 human genes in ExpiCHO was performed using the expression vector pKTH16_dPur, developed in-house (**Supplemental Data** and **Supplemental Figure S1A** and **S1B).** Results from this comparison showed different expression profiles depending on r-protein expressed and expression platform used (**Figure 1A**). Even though purified secreted protein titers varied dramatically between the two platforms for some proteins, no platform provided an overall improved expression profile compared to the other. Instead, each platform provided improved expression in a protein-feature specific manner. Notably, a significant correlation between titer fold changes between the ExpiCHO and QMCF platforms and both protein size (R=-0.5, p=0.015) and glycosylation (R=0.44, p=0.044) was observed (**Figure 1B, Supplemental Figure S1C** and **S1D**). This suggested that each platform provided improved expression in a protein-feature specific manner, where larger and less glycosylated proteins tended to have an expression advantage in the QMCF system and vice versa in case of the ExpiCHO platform. We performed gene expression profiling of the ExpiCHO and CHOEBNALT85 cell lines and noticed comparable transcriptional and secretory pathway activities across the two platforms (**Supplemental Figure S1E**). Interestingly, the stable QMCF system showed significantly higher translational utilization than the ExpiCHO system. The protein-feature specific expression profiles observed between the two CHO expression platforms suggested that neither of the CHO expression hosts or cultivation-protocols provided optimal conditions in a protein-independent manner.

**Figure 1.**
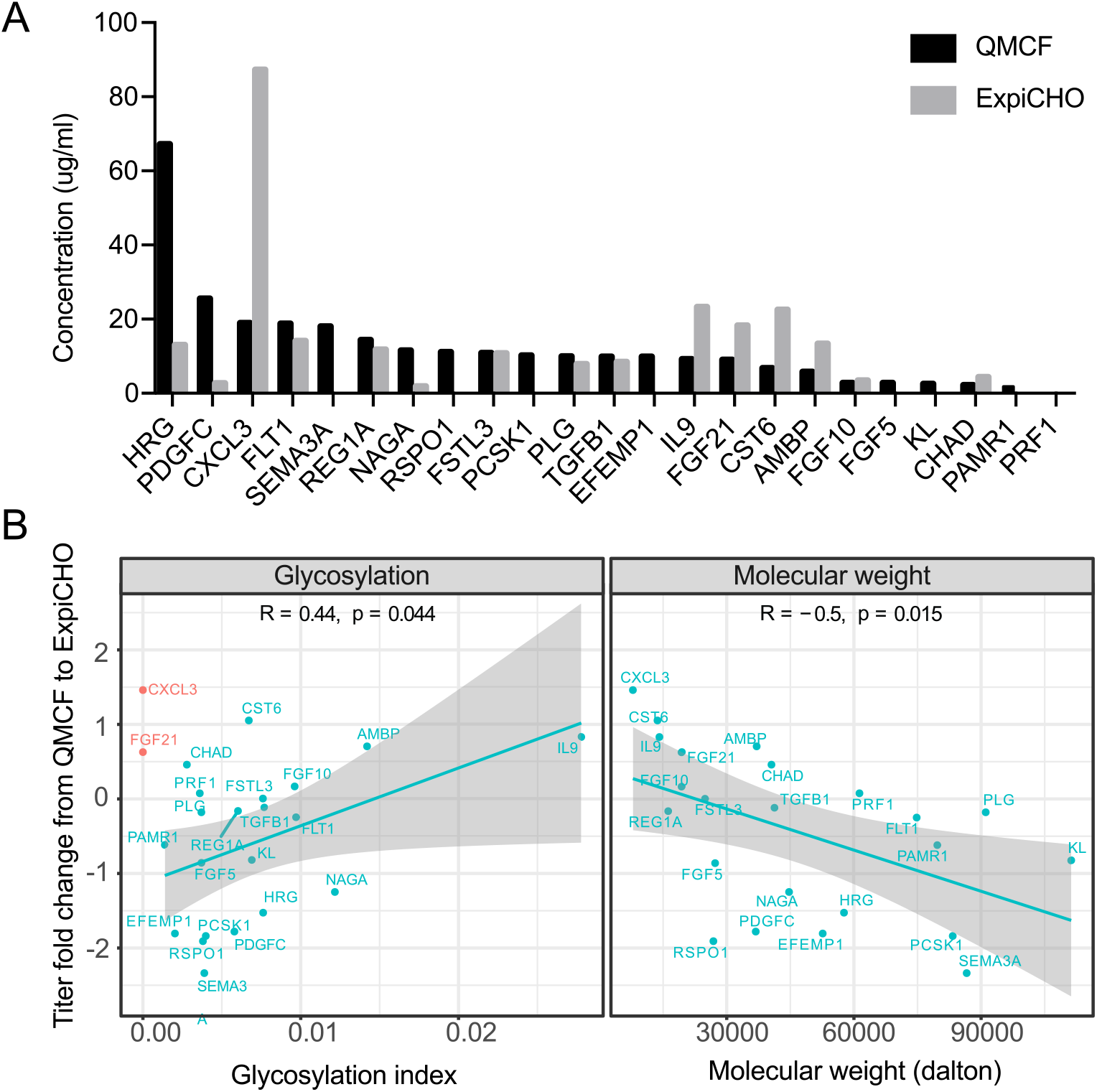
The expression of human secreted proteins in CHO cells. (A) The secreted and purified titers of 23 difficult to express proteins in the QMCF technology versus the ExpiCHO system. (B) Correlation between the molecular weight and r-protein titer fold-change in ExpiCHO compared to the QMCF system. Non-glycosylated proteins are colored red and excluded from the correlation calculation.

### Expression of challenging human proteins in HEK293 resulted in overall improved secreted titers compared to CHO

We hypothesized that a human expression host may have a more compatible secretory pathway for human secreted proteins and hence provide benefits for the expression of those that are particularly difficult in CHO. Thus, we also sought to compare r-protein production between CHO and HEK293. A panel of 24 difficult to express proteins from the Human Secretome Project (Table 1) was expressed side-by-side in CHO and HEK293 cells using a standardized expression- and evaluation-pipeline (**Figure 2A**). We evaluated both an optimized version of the semi-stable QMCF technology, using the CHOEBNALT85-1E9 and 293ALL cell lines, and a fully transient expression setup with 293-F, Freestyle 293-F and Freestyle CHO-S cell lines. In the episomal stable QMCF-system, 9 out of 24 exogenously expressed genes (THBS4, ARTN, BMP10, POSTN, FSTL3, AMBP, CCL28, CXCL13 and NRTN) showed more than 2-fold improved expression in HEK293 (293ALL) compared to CHO cells (CHOEBNALT85-1E9) (**Figure 2B, Supplemental Figure S2A**). On the other hand, only one gene (PLG) showed at least 2-fold improved expression in CHO compared to HEK293. In addition, for six of the genes (NRTN, NRTN pp CXCL13, CCL28, CCL20 and LOX) no detectable secreted expression could be observed in the CHO cell line, whereas 3 genes (CCL20, LOX and PLG) resulted in no detectable protein expression in case of 293ALL.

**Table 1.**
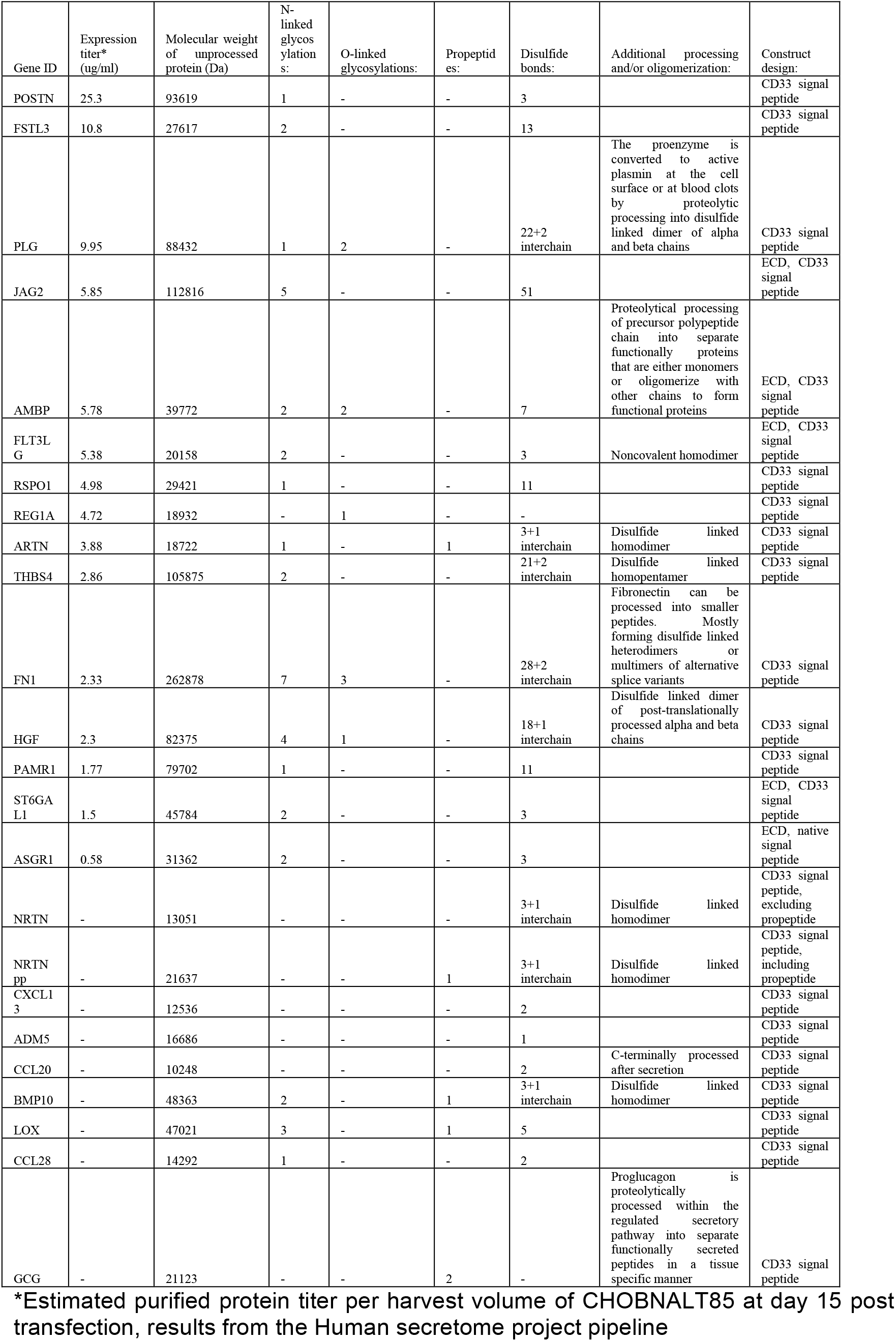
Overview of 24 difficult to express proteins evaluated side-by-side in HEK293 and CHO.

| Gene ID | Expression titer* (ug/ml) | Molecular weight of unprocessed protein (Da) | N-linked glycosylation s: | O-linked glycosylations: | Propeptides: | Disulfide bonds: | Additional processing and/or oligomerization: | Construct design: |
| --- | --- | --- | --- | --- | --- | --- | --- | --- |
| POSTN | 25.3 | 93619 | 1 | - | - | 3 |  | CD33 signal peptide |
| FSTL3 | 10.8 | 27617 | 2 | - | - | 13 |  | CD33 signal peptide |
| PLG | 9.95 | 88432 | 1 | 2 | - | 22+2 interchain | The proenzyme is converted to active plasmin at the cell surface or at blood clots by proteolytic processing into disulfide linked dimer of alpha and beta chains | CD33 signal peptide |
| JAG2 | 5.85 | 112816 | 5 | - | - | 51 |  | ECD, CD33 signal peptide |
| AMBP | 5.78 | 39772 | 2 | 2 | - | 7 | Proteolytical processing of precursor polypeptide chain into separate functionally proteins that are either monomers or oligomerize with other chains to form functional proteins | ECD, CD33 signal peptide |
| FLT3LG | 5.38 | 20158 | 2 | - | - | 3 | Noncovalent homodimer | ECD, CD33 signal peptide |
| RSPO1 | 4.98 | 29421 | 1 | - | - | 11 |  | CD33 signal peptide |
| REG1A | 4.72 | 18932 | - | 1 | - | - |  | CD33 signal peptide |
| ARTN | 3.88 | 18722 | 1 | - | 1 | 3+1 interchain | Disulfide linked homodimer | CD33 signal peptide |
| THBS4 | 2.86 | 105875 | 2 | - | - | 21+2 interchain | Disulfide linked homopentamer | CD33 signal peptide |
| FN1 | 2.33 | 262878 | 7 | 3 | - | 28+2 interchain | Fibronectin can be processed into smaller peptides. Mostly forming disulfide linked heterodimers or multimers of alternative splice variants | CD33 signal peptide |
| HGF | 2.3 | 82375 | 4 | 1 | - | 18+1 interchain | Disulfide linked dimer of post-translationally processed alpha and beta chains | CD33 signal peptide |
| PAMR1 | 1.77 | 79702 | 1 | - | - | 11 |  | CD33 signal peptide |
| ST6GAL1 | 1.5 | 45784 | 2 | - | - | 3 |  | ECD, CD33 signal peptide |
| ASGR1 | 0.58 | 31362 | 2 | - | - | 3 |  | ECD, native signal peptide |
| NRTN | - | 13051 | - | - | - | 3+1 interchain | Disulfide linked homodimer | CD33 signal peptide, excluding propeptide |
| NRTN pp | - | 21637 | - | - | 1 | 3+1 interchain | Disulfide linked homodimer | CD33 signal peptide, including propeptide |
| CXCL13 | - | 12536 | - | - | - | 2 |  | CD33 signal peptide |
| ADM5 | - | 16686 | - | - | - | 1 |  | CD33 signal peptide |
| CCL20 | - | 10248 | - | - | - | 2 | C-terminally processed after secretion | CD33 signal peptide |
| BMP10 | - | 48363 | 2 | - | 1 | 3+1 interchain | Disulfide linked homodimer | CD33 signal peptide |
| LOX | - | 47021 | 3 | - | 1 | 5 |  | CD33 signal peptide |
| CCL28 | - | 14292 | 1 | - | - | 2 |  | CD33 signal peptide |
| GCG | - | 21123 | - | - | 2 | - | Proglucagon is proteolytically processed within the regulated secretory pathway into separate functionally secreted peptides in a tissue specific manner | CD33 signal peptide |

\*Estimated purified protein titer per harvest volume of CHOBNALT85 at day 15 post transfection, results from the Human secretome project pipeline

**Figure 2.**
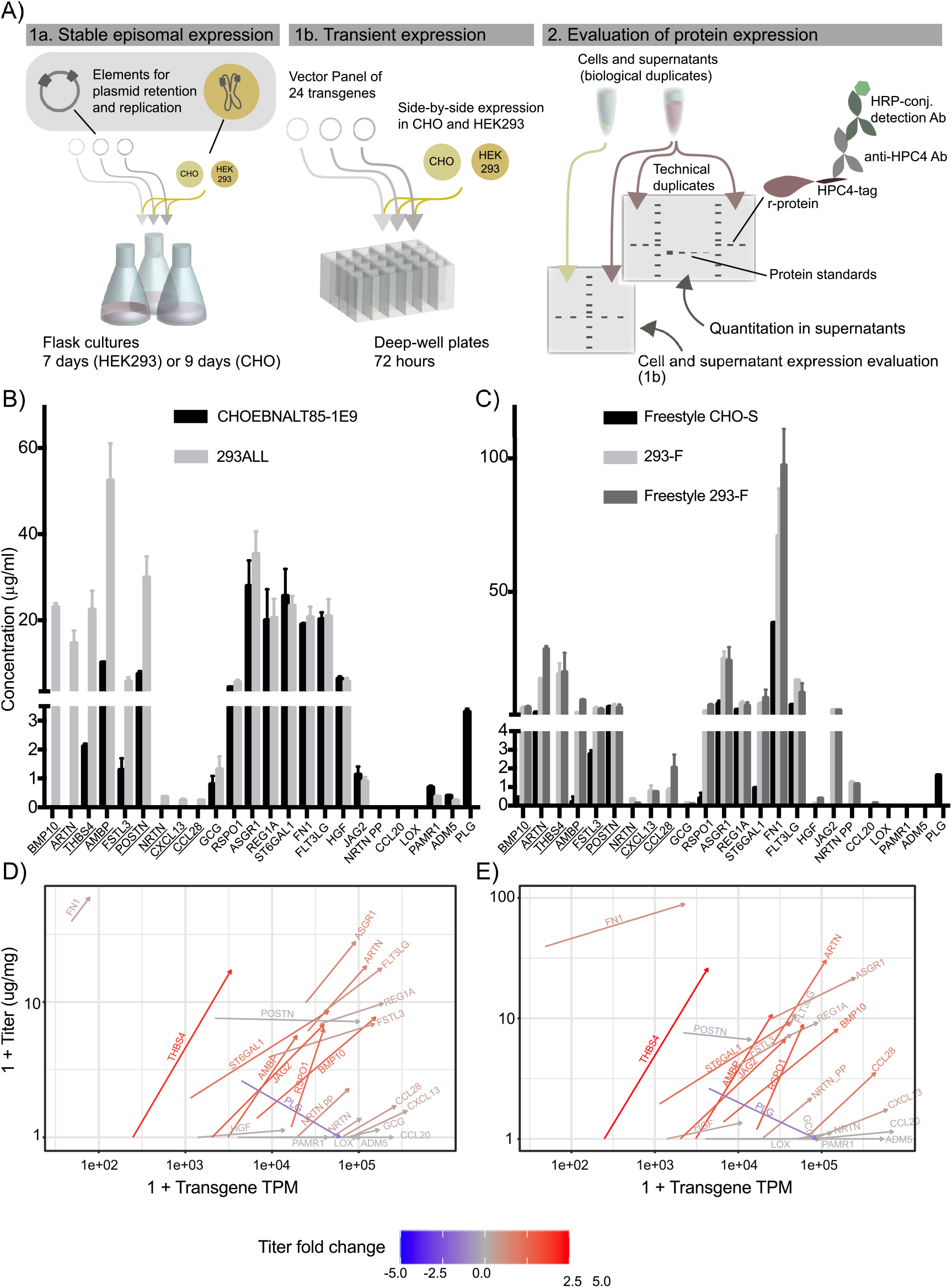
HEK293 provides improved secreted titers of difficult to express proteins compared to CHO in two different expression systems. Comparison of expression titers from supernatants of HEK293 and CHO cell lines from (A) a vector panel of the 24 transgenes was expressed side-by-side in HEK293 and CHO cells in both medium-scale (shake flasks) in the stable episomal long-term expression system QMCF in CHOEBNALT85-1E9 and 293ALL cell lines (1a) and in small-scale (deep-well plates) transient cultivations of Freestyle CHO-S, 293-F and Freestyle 293-F (1b). The protein expression was analyzed by western blot where r-protein was detected by targeting the C-terminal HPC4-tag using an anti-HPC4 antibody. Mean secreted titers ± SD of difficult to express proteins expressed in (B) the stable episomal expression system pQMCF using the CHOEBNALT-85-1E9 and 293ALL cell lines or (C) the transient cultivation protocol of Freestyle CHO, 293-F and Freestyle-293-F. The underlined gene names indicate genes with more than two-fold improved expression in HEK293 cell lines compared to CHO. Differences in protein titer and mRNA levels were quantified for each expressed transgene in CHO and HEK293 cells. The transgene RNA and protein amounts for Freestyle CHO-S and 293-F cultures (D) and Freestyle 293-F (E) cultures are plotted on the X and Y-axis, respectively. For each protein, the changes in transgene and protein levels from CHO to HEK293 are represented by an arrow. The fold changes in protein levels in HEK293 cells are color-coded.

In the small-scale fully transient expression setting the improvement in secreted expression using HEK293 compared to CHO was even more profound. For transient protocol setup and harvest data see **Supplemental data** and **Supplemental Figure S2B** and **S2C**. Protein titer estimations of cell culture supernatants are shown in **Figure 2C** and western blots from all proteins are shown in **Supplemental Figure S2D**. The results showed more than two-fold higher secreted titers of 16 of the 24 difficult to express proteins in both of the closely related HEK293 cell lines compared to the CHO cell line. Only one protein, PLG, showed at least two-fold higher expression in CHO cells compared to HEK293. For half of the evaluated genes (12 of 24), no or only traces of protein could be detected in supernatants from CHO cells. Moreover, for nine of the investigated r-proteins (THBS4, CCL28, CXCL13, CCL20, HGF, PAMR1, LOX, NRTN and NRTN pp) no or only traces of protein could be detected in both the culture supernatant and cell lysate of CHO cells (**Supplemental S2D**). In the case of expression in HEK293 cells, only four genes (PAMR1, LOX, ADM5 and PLG) resulted in no or only traces of secreted protein in both HEK293 cell lines. However, three of these proteins (PAMR1, LOX and ADM5) could be detected in cell lysates of both HEK293 cell lines, suggesting inefficient secretion from HEK293 cells. Strikingly, only one of the 24 investigated proteins, PLG, could not be detected in either cell lysates or cell supernatants in any of the HEK293 cell lines. For a subset of supernatant samples, the relative titer change between cell lines were confirmed using liquid chromatography tandem mass spectrometry (LC-MS/MS) combined with protein quantification based on the SIS PrEST technology (see **Supplemental Data** and **Supplemental Table 1**) (Edfors et al., 2014). Results from the MS/MS analysis showed the same expression trends between HEK293 and CHO cells for each r-protein investigated compared to the western blotting data (**Supplemental Figure S2E**). As the two methodologies in this case depend on different parts of the polypeptide sequences for r-protein detection, depending on r-protein processing in the samples and differences in r-protein processing between cell lines, protein titers of either method reported here should be considered estimates and not absolute quantities.

Taken together, 8 out of 24 genes (BMP10, ARTN, THBS4, AMBP, FSLT3, NRTN, CXCL13 and CCL28) consistently expressed better in HEK293 cells in both the semi-stable and the transient expression comparison. For CHO, the only gene that expressed better compared to HEK293 in both setups was PLG.

### Transcriptome profiling showed variation in secretory pathway utilization between HEK293 and CHO driven by limited set of gene outliers

Based on the observed improvement of the expression of several human r-proteins in HEK293 compared to CHO, a transcriptomic comparison between the two cell lines was performed with emphasis on r-protein transcript levels and host gene expression patterns. Initially, transcript levels for each transgene in the panel were quantified and compared to the r-protein titers in supernatants. Overall, both HEK293 cell lines showed elevated transgene transcript levels compared to CHO cells, consistent with the generally higher transfection efficiency observed for HEK293 compared to CHO (**Figure 2D-E, Supplemental Figure S2F**). However, neither the transgene transcript abundance nor its fold changes correlated with secreted protein titers (**Supplemental Figure S2G**). Notably, some of the proteins with the highest transgene abundances (CCL20, CCL28, CXCL13 and ADM5) displayed among the lowest titers of secreted protein, or no secretion at all. This suggests that such extreme transgene mRNA levels may come at the cost to endogenous gene expression. Alternatively, this could indicate inefficient translation of the transgene mRNAs resulting in an accumulation of untranslated mRNAs in the cell. Moreover, among the proteins with the highest secreted titers in HEK293 (FN1 and THBS4), mRNA levels of the transgene were among the lowest in the data set (between 40 – 4200 TPM).

Furthermore, the mRNA expression levels across major molecular processes in CHO and HEK293 cells were quantified, both for cells expressing the panel of r-proteins (expressing cells) and cells transfected with empty plasmid (non-expressing cells) (**Figure 3A**). Between expressing and non-expressing cells within each cell line, the transcriptome usage were similar for both CHO and HEK293, with the exception of transcription of genes related to translation and signaling molecules in the Freestyle 293-F cell line. Overall, HEK293 and CHO cells showed great variation in the utilization of their respective transcriptomes. Most evidently, genes associated with translation showed lower mRNA expression on average in CHO cells, compared to HEK293 cell lines, whether a transgene is being expressed or not. In addition, the CHO cells showed a higher proportion of their transcriptome expression focused on biosynthesis, central carbon metabolism and signal transduction. Interestingly, HEK293 cells showed an overall less active secretory pathway compared to CHO (**Figure 3A** and **Supplemental Figure 3A**), despite secreting higher amounts of r-proteins. However, the overall gene expression levels were generally higher in HEK293 compared to CHO of all secretory pathway subgroups, with the exception of protein folding that was significantly higher in CHO (**Figure 3B**). Instead, the higher fraction of the transcriptome devoted to the secretory pathway in CHO cells compared to HEK293 was due to a very small subset of highly expressed secretory pathway genes whereas HEK293 cells, on the other hand, expressed a more diverse set of secretory pathway genes (**Supplemental Figure S3B**). While the fraction of the transcriptome, across all samples, that was utilized for secretory pathway genes did not correlate with transgene expression levels nor estimated secreted r-protein titers (**Supplemental Figure S3C**), there was a linear relationship between expression of several secretory pathway genes and transgene mRNA abundances in CHO producers (**Supplemental Figure S3D).** In HEK293 cells however, peak secretory pathway activities occur in clones with low to medium transgene load. This suggested a saturation of the secretory pathway in HEK293 cells, which may be a result of the exceptionally high transgene mRNA loads observed in case of several transgenes. Across all samples, a significant negative correlation was observed between r-protein titer and gene expression within the protein folding and ER glycosylation functional groups, respectively (**Supplemental Figure S3E**). Focusing on individual secretory pathway genes, overall similar expression levels were observed between the two cell lines with the exception of a limited set of extreme gene outliers (**Figure 3C**). Amongst these outliers, there was a substantial group of genes with very low or no expression in CHO whereas the expression in HEK293 was moderate to high (**Figure 4A**). Within this group of genes we observed several genes with previous support of impact on r-protein secreted titers (Table 2)(Le Fourn et al., 2014; Hansen et al., 2015; Haredy et al., 2013; Hwang et al., 2003; Ishaque et al., 2007; Lasunskaia et al., 2003; Lee et al., 2009; Meleady et al., 2011; Ohya et al., 2008; Orellana et al., 2018; Sommeregger et al., 2016).

**Table 2.** Overview of differentially expressed genes between HEK293 and CHO evaluated based on their impact on productivity.

| Gene ID: | log2 Fold Change: | p.adj.: | Secretory pathway subgroup: | Previous association with protein productivity: |
| --- | --- | --- | --- | --- |
| HSPA1B | 12.89 | 0 | ERAD | Previous data support 1) enhanced yields of hybridomas generated with Hsp70-expressing mouse myeloma NS0 cells compared to wt cells (Lasunskaja et al., 2003); 2) increased yields of rFVIII in BHK-21 cells co-expressing Hsp70 (Ishaque et al., 2007); 3) increased culture longevity and IFN-gamma titers of CHO cells coexpressing Hsp70 (Lee et al., 2009). |
| EIF2AK2 | 12.71 | 1.8E-213 | UPR | Previous study showed up-regulated expression of EIF2AK2 in HP antibody-producing CHO clone compared to LP (Orellana et al., 2018). |
| TBC1D9 | 12.32 | 2.0E-206 | Trafficking |  |
| HSPA4L | 12.07 | 0.0E+00 | ERAD |  |
| RAB11FIP1 | 11.86 | 5.2E-192 | Trafficking |  |
| GALNT18 | 9.98 | 1.2E-188 | Golgi glycosylation |  |
| SNAP25 | 9.48 | 2.9E-117 | Trafficking |  |
| MGAT3 | 9.28 | 1.0E-117 | Golgi glycosylation |  |
| MYO5B | 9.17 | 3.1E-211 | Trafficking |  |
| AGAP2 | 8.05 | 9.2E-92 | Trafficking |  |
| SVIP | 7.07 | 2.1E-67 | ERAD |  |
| RAB6B | 6.33 | 7.0E-98 | Trafficking |  |
| DERL3 | 5.62 | 1.0E-254 | ERAD |  |
| JUN | 3.03 | 0 | Trafficking | Previous study showed up-regulated expression of JUN in HP antibody-producing CHO clone compared to LP (Orellana et al., 2018). |
| SRP9 | 2.85 | 0 | Translocation | In a study by Le Fourn et al. (2014), expression of SRP9 along with other components of the SRP complex increased antibody expression in CHO |
| PDIA4 | 1.78 | 1.6E-24 | Protein folding | PDIA4 was found expressed at higher levels in CHO cells producing easy to express scFv compared to CHO cells producing difficult to express scFv (Sommeregger et al., 2016). |
| ATF4 | 1.71 | 1.3E-134 | UPR | Overexpression of ATF4 has previously been reported to improve the productivity of antithrombin III and IgG, respectively, in CHO cells (Ohya et al., 2008; Haredy et al., 2013). |
| RAB31 | -3.18 | 0 | Trafficking |  |
| RAB43 | -2.88 | 2.0E-167 | Trafficking |  |
| PDIA3 | -2.52 | 0 | Protein folding | Shown to be expressed at higher levels in CHO cells producing easy to express scFv variant than CHO cells producing difficult to express scFv (Sommeregger et al 2016). Conflicting effects on productivity have been reported, with both positive (Hwang et al., 2003) and negative impacts (Hansen et al., 2015). |
| HSPA8* | -2.42 | 0 | Protein folding | Upregulation of HSPA8 correlated with sustained productivity of antibodies in CHO cells over 10 days fed-batch cultures (Meleady et al., 2011). |

Abbreviations: LP: Low producer; HP: High producer

**Figure 3.**
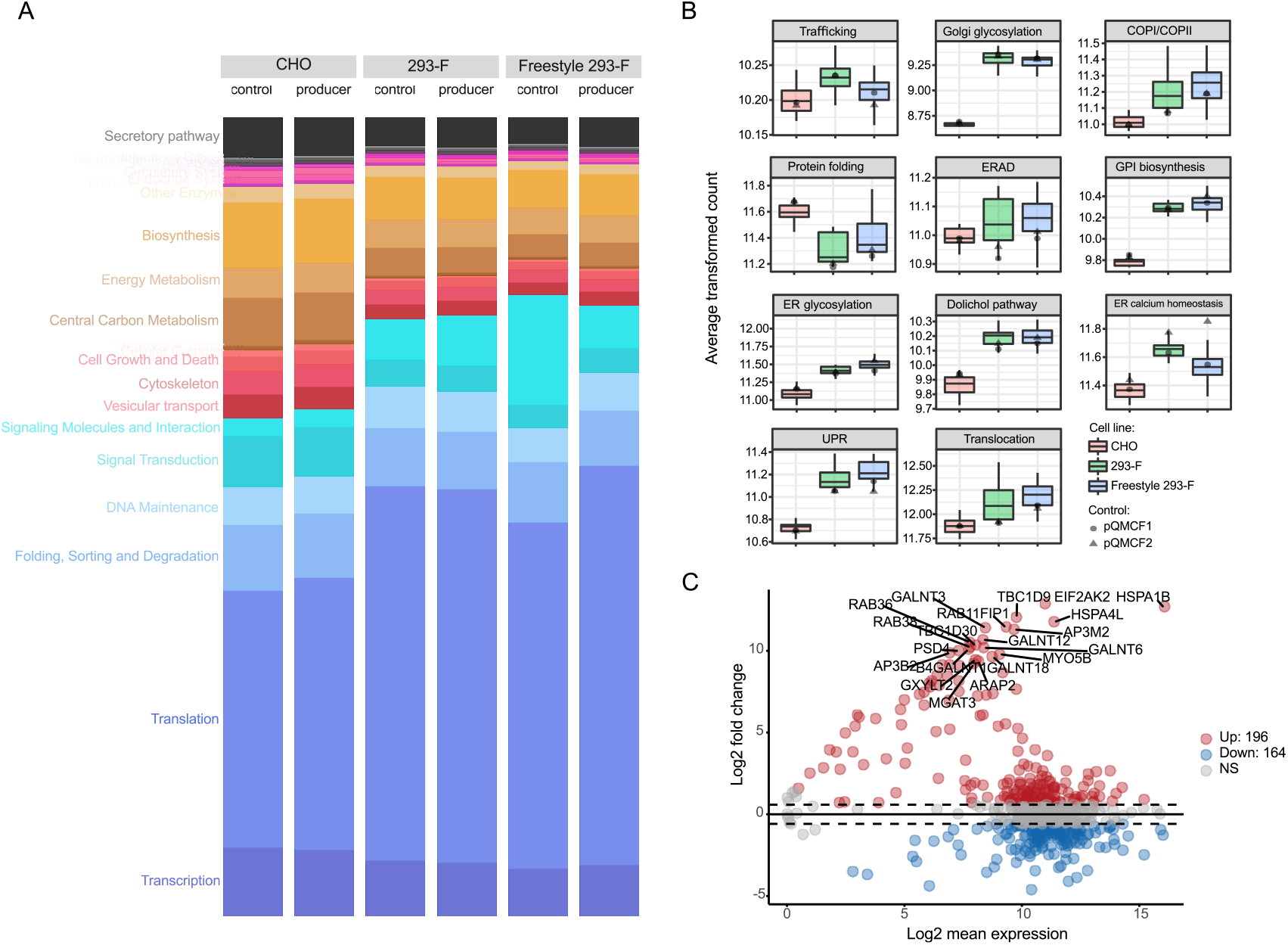
The utilization of the secretory pathway is distinctly different between CHO and HEK293 as a result of a limited set of gene outliers. (A) Comparison of overall transcriptome usage across cell lines and producers. The fraction of the transcriptome dedicated to various cellular functions is represented by the height of each bar, where related pathways are colored similarly. The transgene mRNA levels were excluded. (B) Overall gene expression of 11 functional secretory pathway subgroups in CHO and HEK293 upon overexpression of target genes. For each group, the average gene expression level for the sample was computed and plotted for all samples. Black dots represent average expression in each secretory pathway group for control samples with an empty vector (pQMCF-plasmid). (C) Differential expression MA-plot showing the mean expression and fold-changes for the secretory pathway genes between HEK293 and CHO. Positive fold-changes denote higher expression in HEK293 and vice versa. The top 20 most differentially expressed secretory pathway components are labeled.

**Figure 4.**
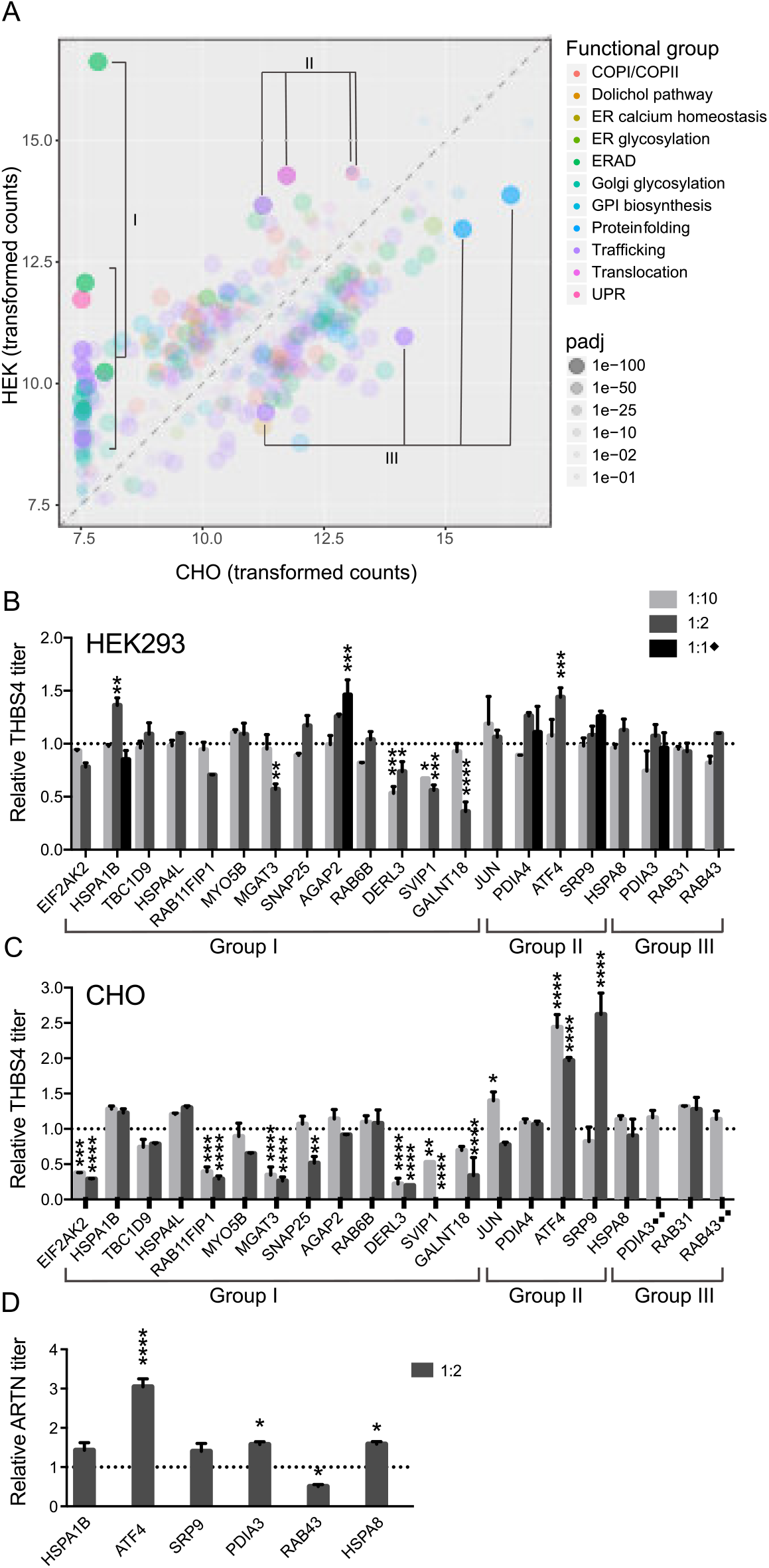
Evaluation of outlier secretory pathway genes on secreted protein titers. (A) Comparison of expression of individual secretory pathway genes, with transformed counts for CHO and HEK293 shown on the X-and Y-axis respectively. Most secretory pathway genes lie within the shaded region drawn around the identity line (x=y), showing an overall conserved pattern of expression. Differentially expressed secretory pathway genes evaluated based on effects of the expression of a difficult-to-express protein are highlighted and divided into three groups (I, II and III) based on expression levels in the two cell lines. All other genes are shaded. Mean relative secreted titers ± SD (N = 2) of THBS4 determined by ELISA in HEK293 (B) and CHO (C) when co-expressed with differentially expressed secretory pathway genes (x axis) compared to the expression level of THBS4 alone (empty vector). Plasmid ratios of 1:1, 1:2 or 1:10 (secretory pathway gene:THBS4) upon transfection was evaluated. ♦The 1:1 plasmid ratio was only evaluated for HSPA1B, AGAP2, ATF4, SRP9 and PDIA3 in HEK293 cells. ♦♦Only the 1:10 plasmid ratio evaluated were evaluated for PDIA3 and RAB43 in CHO cells. D) Co-expression results of ARTN expression levels (determined by western blot) when combined with a subset of the differentially expressed secretory pathway genes in B and C in CHO with plasmid ratios 1:2 (secretory pathway gene:ARTN). Mean relative ARTN titers ± SD (N = 2) between supernatants of cells co-transfected with secretory pathway genes versus no transgene (empty vector). Significant different expression of THBS4 (B and C) or ARTN (D) compared to the co-expression with an empty vector control (determined by one-way ANOVA and Dunnett’s test) are indicated by the asterisk sign (* P adj ≤ 0.05; ** P adj ≤ 0.01; *** P adj ≤ 0.001; **** P adj ≤ 0.0001).

### Highly expressed helper proteins in HEK293 compared to CHO have a positive impact on secretion of difficult to express proteins when co-expressed also in CHO cells

We hypothesized that the secretory pathway genes with extreme expression variation between the cell lines may contribute to profound differences in productivity observed between the cell lines for some human difficult to express proteins. Consequently, a set of secretory pathway genes was co-expressed with the difficult to express protein THBS4 in CHO and HEK293 to evaluate their impact on secreted productivities. The selection of gene outliers to evaluate was based on the highest differential expression between the cell lines but also on previous literature supporting potential roles in protein secretion or demonstrated effects on bioproductivity (**Table 2**; **Supplemental Table S4**) (Le Fourn et al., 2014; Hansen et al., 2015; Haredy et al., 2013; Hwang et al., 2003; Ishaque et al., 2007; Lasunskaia et al., 2003; Lee et al., 2009; Meleady et al., 2011; Ohya et al., 2008; Orellana et al., 2018; Sommeregger et al., 2016). The selected genes were divided into three groups (I, II and III) based on expression levels in the two cell lines (**Figure 4A**). Three genes (HSPA1B, AGAP2 and ATF4) had a small, albeit significant, positive effect on the secreted titer of THBS4 compared to cells only expressing THBS4 (**Figure 4B**). For CHO cells a more profound titer improvements was observed with genes in group II (**Figure 4C**), including SRP9, ATF4 and JUN that had moderate expression endogenous expression in CHO but higher expression in HEK293. More than two-fold improvements in secreted THBS4 titers were observed in CHO when co-expressed with ATF4 or SRP9, and a 1.5-fold titer increase was observed when co-expressed with JUN. In addition, slight improvements (however not statistically significant) of THBS4 titers in CHO cells were observed when co-expressed with some secretory pathway genes from groups I and III such as HSPA1B, HSPA4L and RAB31, depending on the plasmid ratio between the co-expressed transgenes. Validation of a subset of the gene outliers in combination with ARTN in CHO cells (**Figure 4D**) showed a significant positive impact of ATF4, PDIA3 and HSPA8 on ARTN secretion. Moreover, HSPA1B and SRP9 overexpression generated a small increase, albeit not significant, in ARTN titers. Higher secreted ARTN titers, but not THBS4, in CHO cells were associated with lower viable cell densities and viability at harvest compared to controls (**Supplemental Figure S4**). This may relate to differential effects of the two r-proteins on cells. In addition, several outlier secretory pathway genes of group I (EIF2AK2, RAB11FIP1, MGAT3, DERL3, SVIP1 and GALNT18), with low or no expression in CHO, but moderate to high expression in HEK293, significantly decreased secreted THBS4 titers dramatically when added exogenously in CHO. A similar trend was observed for some of these genes upon co-expression in HEK293 even though the negative impact on secreted THBS4 titers was not as profound. In the case of some of these co-expressed genes (DERL3, MGAT3 and EIF2AK2), these effects were likely the result of a negative impact on cell growth in CHO but not in HEK293, suggesting that these genes are not compatible with the CHO cell machinery of maintaining cellular growth and productivity.

Since both THBS4 and ARTN were profoundly better expressed in HEK293 compared to CHO and the low expression of these proteins in CHO could be rescued by the overexpression of secretory pathway components expressed at higher levels in HEK293, we hypothesize that such differences in secretory pathway components may be beneficial for the secretion of difficult to express proteins in HEK293.

### Proteasomal and propeptide convertase genes were differentially expressed between CHO and HEK293

To obtain a more fine-grained understanding of the cell line differences, differential expression analysis comparing HEK293 to CHO cells was performed. Due to drastic organismal differences, many genes outside of the secretion machinery showed distinct expression between CHO and HEK293 cells (**Supplemental Figure S5A, Supplemental Table S4**). In fact, more than 80% of the genes were significantly differentially expressed between the two organisms. Few canonical pathways were consistently up- or down-regulated in one cell line compared to another, as shown by gene-set enrichment analysis (**Supplemental Figure S5B**). However, among them the proteasome protein family, which degrades misfolded proteins in a controlled fashion, was expressed at significantly higher levels in CHO compared to HEK293 cells (**Supplemental Figure S5C**). Another protein family that showed significantly different expression profiles between HEK293 and CHO is the propeptide convertase family (**Supplemental Figure S5D**). HEK293 showed significant upregulation of PCSK2, PCSK4, PSCK5, PCSK6 and PCSK8 compared to CHO. On the other hand, CHO expressed higher levels of PCSK1, PCSK3 and PCSK7.

### Differentially activated secretory pathway genes between HEK293 and CHO upon transgene expression

To evaluate overall differences in cellular dynamics between HEK293 and CHO cells when producing r-proteins and to better account for organismal disparity, we calculated the average activation of genes by determining the differential expression upon transgene expression across cell lines during r-protein production with non-producer cells as reference (**Figure 5A**). While the majority of the genes were not differentially activated in either of the cell lines, several genes showed significantly opposite trends in HEK293 and CHO cells (**Supplemental Table S5**). The top 20 genes identified as differentially activated in producers between CHO and HEK293 strongly enriched for members of the secretory pathway (hypergeometric p-value < 0.0005). Among them, four genes (SEC61A1, SEC63, DNAJC3 and ERO1B) are directly involved in posttranslational functions of the secretory pathway. Co-expression evaluation of SEC61A1, DNAJC3 or ERO1B between HEK293 and CHO, did however not result in improved secreted expression of THBS4 in either cell line (**Supplemental Figure S6**). On the contrary, overexpression of all these genes had a significant detrimental effect on secreted titers in CHO cells. This may suggest that the activation of these genes is likely a result of, rather than the cause of a systemic response upon transgene expression and that further activation of these genes has no, or detrimental, effects on protein production.

**Figure 5.**
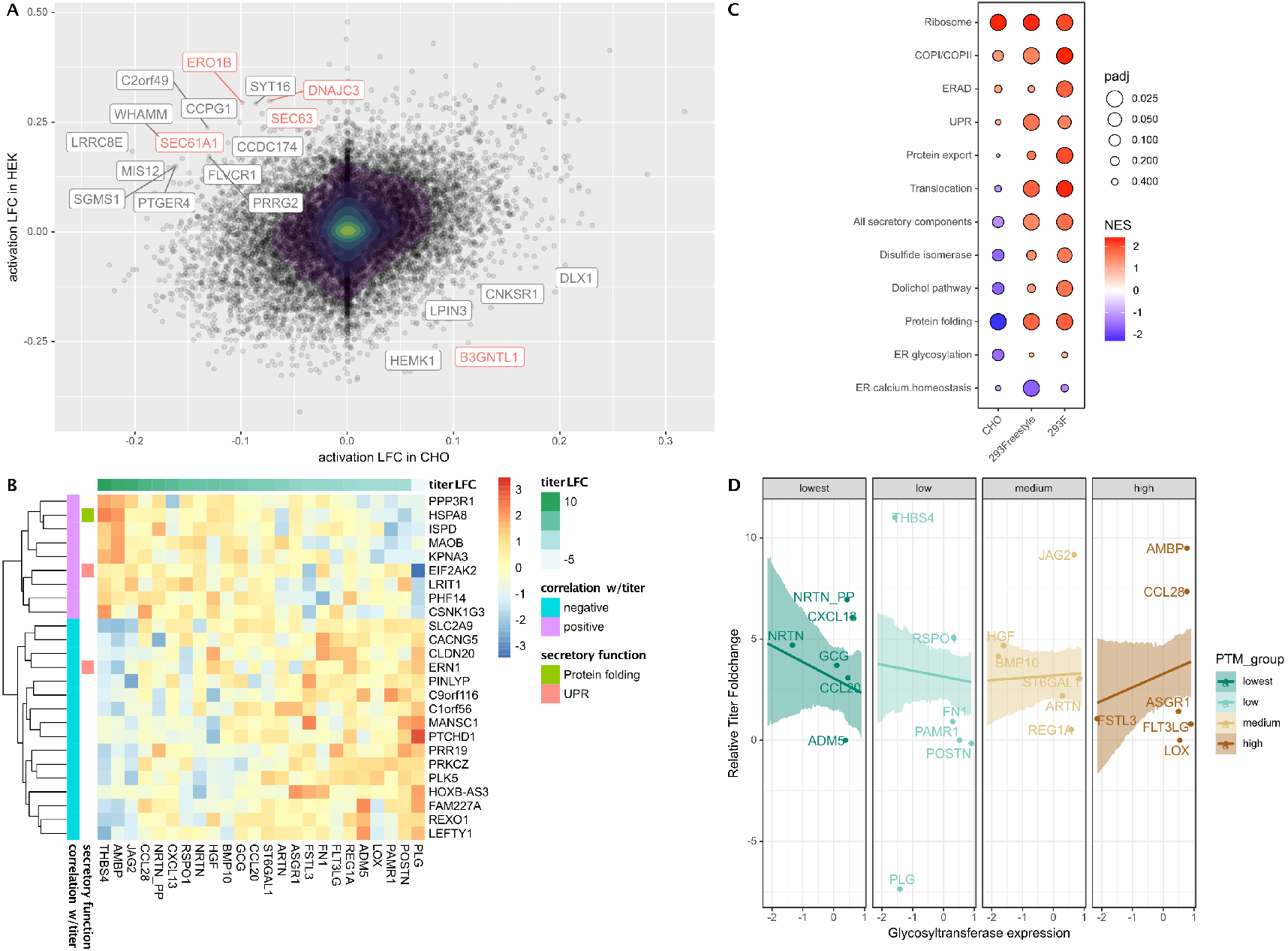
Transgene expression induces differential activation of genes between cell lines and between different r-proteins. (A) Comparison of gene activation upon transgene expression between HEK293 and CHO. Log2 fold-changes (LFCs) are calculated based on the differential expression between producers and controls for CHO and HEK293 cells, and are plotted on the X- and Y-axis respectively. Genes showing the most divergent activation patterns between HEK293 and CHO are labeled. The top differentially activated genes are enriched for secretory pathway genes (colored red, hypergeometric p-value = 0.004). (B) Heatmap of endogenous secretory pathway genes with the highest positive or negative correlation between differential activation and r-protein titer change from CHO to HEK293. In each entry, a more positive fold-change of the endogenous secretory pathway gene (Y-axis label) for a given r-protein (X-axis label) indicates stronger activation in HEK293 compared to CHO when producing that protein. (C) Overall activation of secretory pathway subsystems. The dots in the scatter heatmap show pathway activities (y-axis) across cell lines (x-axis), with colors indicating activation/suppression and sizes indicating the corresponding significance. (D) Differences in activation of O- and N-linked glycosyltransferases between HEK293 and CHO cells correlate with titer improvement of moderately-to heavily-glycosylated r-proteins. R-proteins harbouring more frequent N- and O-glycosylation sites per AA residue (rightmost panel) tend to benefit more from increased expression of glycosyltransferases from CHO to HEK, while a reversed trend was observed for non- and lightly-glycosylated r-proteins (leftmost panel).

To see pathways that were preferentially activated in each cell line, gene set enrichment analysis (GSEA) was performed using the fold changes of differential activation for each cell line (**Figure 5C**). The producers in all three cell lines significantly upregulated genes involved in translation. However, HEK293 cells showed higher expression for genes related to protein secretion, compared to CHO cells. For example, translocation and protein export were much more strongly activated in both 293-F and Freestyle 293-F. Genes involved in protein folding such as molecular chaperones, were significantly downregulated in CHO cells, while significantly upregulated in the HEK293 cell lines. Similar pathway activation was observed between the two HEK293 cell lines variants, with the only notable exception being proteasomal function, whose activations were more strongly in Freestyle 293-F than in 293-F.

To systematically identify genes with potential impact on productivity of the cell lines, we calculated for each gene the correlation between its differential activation (**Supplemental Table S6)** and the r-protein titer changes from HEK293 to CHO across clones. Overall, only a small number of the genes displayed r-protein titer change-dependent differential activation. The top genes with the highest positive or negative correlation are given in **Figure 5B**. Results showed that three secretory pathway genes (EIF2AK2, HSPA8 and ERN1) correlated in titer and activation change between HEK293 and CHO. Interestingly, HSPA8 and EIF2AK2 were also found amongst the genes with high expression fold-change between HEK293 and CHO cells (**Supplemental Table S4**).

Beyond the differences in titer improvements, the r-proteins are diverse in their PTM compositions (**Table 1**), utilizing distinct sets of enzymes. With this, we explored whether the differential expression of the enzymes responsible for some of the PTMs between CHO and HEK293 can explain the variation in titer improvements seen by different r-proteins. We posited that proteins with more frequent PTM sites may be more sensitive to changes in the expression of the enzyme responsible for the PTM in question. To quantify the degrees to which certain PTMs are overrepresented in each r-protein, we calculated a “PTM-index” for each r-protein - PTM combination based on the PTM site densities in each r-protein (**Supplemental Figure 7**). Among the three most ubiquitous PTMs taking place within the secretory pathway-disulfide bond, GPI anchor and N-/ O-linked glycosylation, we saw significant interaction between glycosylation-index and enzyme expression in determining titer improvement. More specifically, the titer improvement for more heavily glycosylated proteins showed a strong positive correlation with the differential expression of glycosyltransferases, whereas for non-and lightly-glycosylated r-proteins, the changes in titer were negatively correlated with the glycosyltransferases fold change (**Figure 5D**).

## Discussion

The CHO cell line is a well-established bioproduction host with readily available expression protocols both in transient and stable settings. However, with the increasing number of new biologics approaching the market, including next-generation biologics such as engineered scaffold proteins and antibody-fusion proteins, the pharmaceutical industry faces new challenges for efficient protein production. But even natural human proteins can pose challenges for bioproduction both in transient and stable expression systems and require laborious process optimization. Due to the clonal divergence of different immortalized cell lines (Feichtinger et al., 2016; Lin et al., 2014; Malm et al., 2020; Stepanenko and Dmitrenko, 2015; Vcelar et al., 2018; Wurm, 2013), the expression of r-proteins may vary considerably between hosts even of the same origin. Indeed, this was observed in this study, where the two CHO-S based expression platforms, QMCF and ExpiCHO, showed protein-dependent differences in secreted titers. The higher secreted titers of smaller r-proteins and/or more heavily glycosylated r-proteins observed in ExpiCHO compared to the QMCF platform (**Figure 1C**) may relate to the high cell densities of cultivation and high amount of plasmid DNA added upon transfection in the ExpiCHO system. We speculate that this may put growth pressure on the cells, making this platform better suited for producing smaller proteins. On the contrary, the lower cell densities of the QMCF system in combination with the reduced cultivation temperature (30°C instead of 37°C) and longer cultivation times may result in lower cellular stress compared to the ExpiCHO system, enabling production of larger proteins. Changes in titer between the platforms could also be related to clonal differences between cell lines, even though the overall transcriptome utilization is comparable between the CHOEBNALT85 and ExpiCHO cell lines (**Supplemental Figure S1E** and **S1F**). Thus, the data suggest that each system can provide protein-specific advantages possibly related to platform differences. On the other hand, a r-protein-independent improved secreted expression of challenging human proteins was observed when changing expression host from CHO to HEK293 (**Figure 2**). One third of the proteins were expressed at more than 2-fold higher titers in HEK293 compared to CHO in both systems.

Overall, a part of the increased expression in transient cultivations of HEK293 can be explained by an increased transfection efficiency and plasmid uptake compared to CHO reflected in the overall higher mRNA abundances observed in HEK293 (**Figure 2, Supplemental Figure S2F)**, in line with previous observations that HEK293 tend to perform well as a transient expression host and are easier to transfect compared to CHO (Jäger et al., 2015; Pham et al., 2006). However, for some r-proteins extremely high transcript levels resulted in very low or no secreted product, suggesting that such extreme transgene mRNA levels could come at the cost to endogenous gene expression. Moreover, since one third of r-proteins showed increased titers in the HEK293 cell lines compared to CHO also in the semi-stable expression system of QMCF, which provides comparable transfection efficiencies between HEK293 and CHO cells (personal communication with Icosagen), we argue that a subset of human difficult to express proteins that consistently express better in HEK293 compared to CHO in both transient and episomal stable expression settings were likely also a result of differences related to the secretory machinery of the two cell lines.

Transcriptomic analysis of transiently expressing HEK293 and CHO cells showed a profound difference in the overall utilization of the transcriptomes between CHO and HEK293 (**Figure 3A**), which is expected due to the different origins of the cell lines. The higher translational machinery activities in HEK293 cells may afford them increased capacity for translating mRNA, although the level of transgene mRNAs seen in this study should not saturate the ribosomal capacity, as transgene with mRNA levels ranging upwards of 20% of total mRNA content has been shown to translate efficiently (Kallehauge et al., 2017). As the secretory pathway is a major determinant of the titers of secreted proteins (Gutierrez et al., 2020), comparisons between the cell lines focused on secretory pathway components. Higher activities were observed in the protein quality control pathways of UPR and ERAD in HEK293 cells compared to CHO (**Figure 3B**), which may impact protein secretion as upregulated transcription of genes associated with these pathways can increase secretory capacities of host cells (Hussain et al., 2014; Prashad and Mehra, 2015). Moreover, genes involved in protein folding were more highly expressed in CHO cells compared to HEK293. Notably, protein folding showed a significant association with decreased protein titer (FDR p-value = 0.0023, **Supplementary Figure S3E**). This could be a cellular response to increased difficult-to-express protein load, especially if the native machinery for r-protein folding is lacking. At the gene level, most secretory pathway genes have similar expression between the two cell lines (**Figure 3C and 4A**). However, a limited set of genes showed extreme variation in expression between the cell lines. For instance, CHO cells do not express many of the moderately expressed secretory machinery genes expressed in HEK293 cells. Since the r-proteins in our screen are all human proteins, it is possible that the lack of compatible secretory components forced the CHO cells to utilize a smaller subset of more generic machinery components, and this lack of specialization could possibly impact secreted titers. Alternatively, the absence of expression of such genes in CHO may be compensated by the expression of other genes without a human ortholog and hence not included in our analysis. Notably, amongst the genes more highly expressed in HEK293 compared to CHO, we identified several examples of genes that have previously been associated with improved protein production (**Table 2**). For instance, both ATF4 and SRP9 have previously been associated with improved recombinant expression in CHO cells either alone or in combination with other ER components (Le Fourn et al., 2014; Haredy et al., 2013; Ohya et al., 2008). In this study we could show a profoundly positive effect of ATF4 also on secreted production of two difficult to express proteins in CHO cells (**Figure 4**), suggesting that this protein act in a more universal way of improving yield. Overexpression of SRP9 significantly increased THBS4 titers dramatically and had a slightly positive, but not statistically verified, impact on ARTN secretion. The already high endogenous expression of these two genes in HEK293 compared to CHO may explain why the positive effect on secreted titers was not as profound in this cell line and moreover, we hypothesize that such differences in secretory pathway components may be beneficial for the secretion of difficult to express proteins in HEK293. In addition, several genes with more moderate impact on the secreted expression of r-proteins in either CHO or HEK293 were identified in this study, supporting the usage of transcriptomic data to shine light upon secretory pathway differences that impact bioproductivity between cell lines. In HEK293 cells, AGAP2, HSPA1B and ATF4 significantly boosted THBS4 secretion, whereas in CHO cells, SRP9, JUN, PDIA3 and HSPA8 had significant positive impact on secretion of either THBS4 or ARTN. As summarized in **Figure 6**, such genes could indicate productivity bottlenecks in CHO cells when expressing a difficult to express protein.

**Figure 6.**
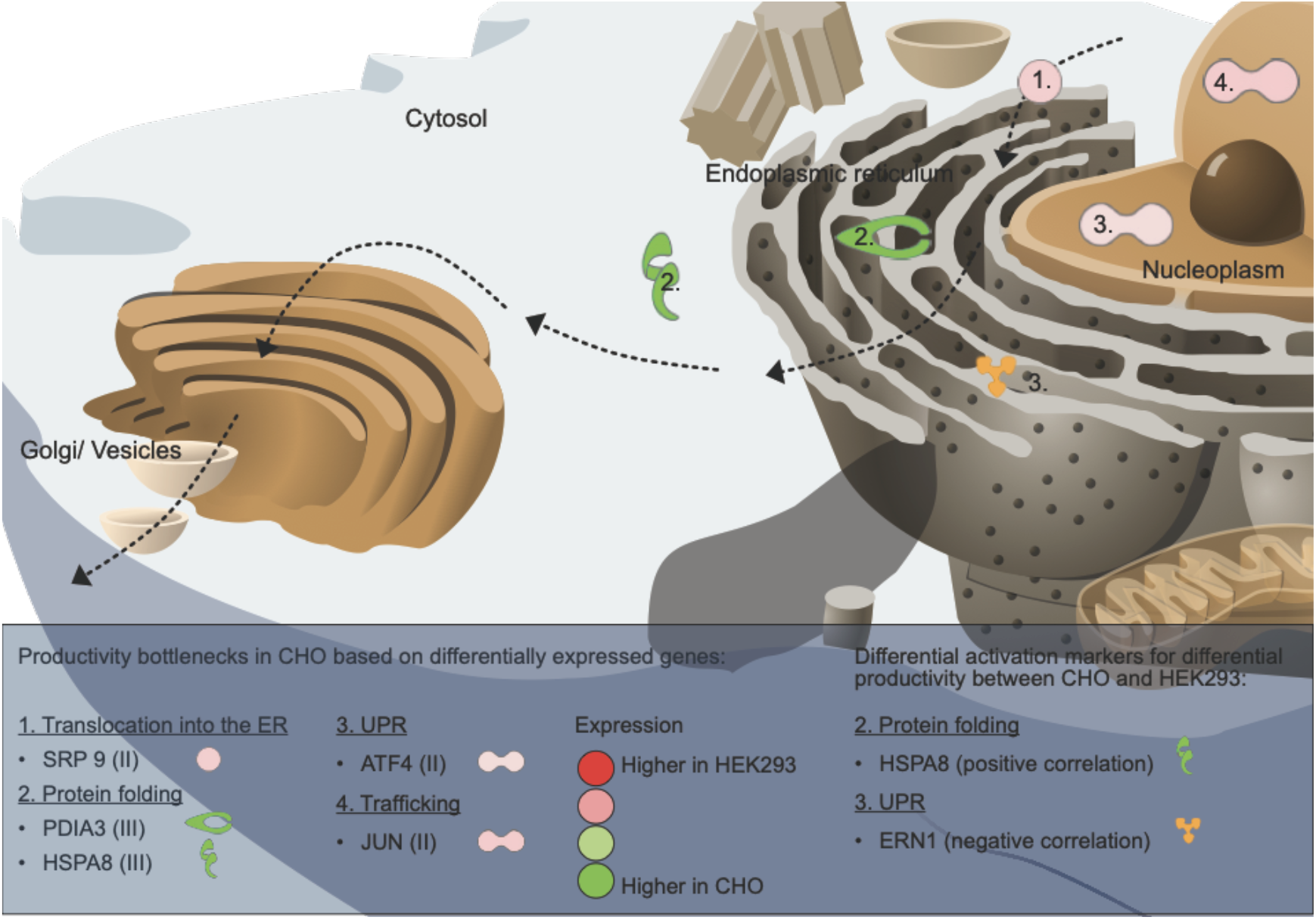
Overview of secretory pathway components with significantly positive impact on productivity in CHO and differential activation markers correlating with differential productivity between CHO and HEK293. The arrows indicate the secretion path of secretory proteins in the cell. Differentially expressed genes between CHO and HEK293 with significantly positive impact (P adj. ≤ 0.05) on THBS4 and/or ARTN titers upon overexpression in CHO are indicated as productivity bottlenecks in CHO cells. The color scale from red to green indicates gene expression fold-change between HEK293 and CHO, where genes of similar expression change between the two cell lines are also grouped into one of three groups (indicated by the roman numerals I, II and III, further described in Figure 4). Differentially activated secretory pathway genes correlating with titer fold-changes between HEK293 and CHO serve as activation markers for titer fold-change of difficult to express protein between cell lines. HSPA8 showed a positive correlation between differential activation upon transgene expression in HEK293 vs. CHO and titer-fold change in HEK293 vs. CHO, whereas ERN1 had a negative correlation. All gene product symbols are mapped into their respective cellular compartments and the numbers indicate the secretory pathway subgroup of these gene products.

Among the top 20 most differentially activated genes, four ER-associated genes (SEC63, SEC61A1, DNAJC3 and ERO1B) were found upregulated in HEK293 but not in CHO upon transgene expression (**Figure 5A**). Even though SEC61A1 along with the other subunits of the translocon has previously been shown to improve the specific productivity of a difficult-to-express antibody in CHO cells (Le Fourn et al., 2014), the individual overexpression of ERO1B, DNAJC3 or SEC61A1 along with THBS4 showed a significantly detrimental effect on productivity in CHO cells in this study (**Supplemental Figure S6**). This may support a less active role of these proteins in the CHO secretory pathway. On the other hand, the significant upregulation of these genes in HEK293 may be well tuned by the cells without further improvements added by exogenous overexpression. Interestingly, differential activation between CHO and HEK293 were overall independent of protein identity or r-protein titer change between the cell lines. This observation suggested that the differential activation upon r-protein expression was mainly driven by cell line differences rather than protein identity or load and highlights cell-line dependent variation in utilization of cellular machinery upon protein production. However, a subset of differentially activated genes correlated in activation fold-change and titer change between the cell lines (**Figure 5B**). Three of these (EIF2AK2, HSPA8 and ERN1) are part of the secretory machinery, out of which HSPA8 has previously been recognized as a marker for protein productivity in CHO cells (Meleady et al., 2011). Notably, EIF2AK2 and HSPA8 were also recognized as extremely differentially expressed between CHO and HEK293. Overexpression of HSPA8 had a slightly positive effect on THBS4 secretion in HEK293 and CHO and a significantly positive effect on ARTN secretion from CHO cells. It’s worth noting that HSPA8 showed stronger activation in HEK293 compared to CHO in THBS4 producing clones, and a reversed activation trend was observed in ARTN producing clones (**Figure 5B**). Hence, HSPA8 expression could be part of the secretory machinery orchestrating differences in protein secretion between the cell lines. On the other hand overexpression of EIF2AK2 showed a negative effect on THBS4 secretion, especially in CHO cells where endogenous EIF2AK2 levels were much lower compared to HEK293. Notably, the correlation between EIF2AK2 activation and titer fold-change between the cell lines was mainly driven by the strong negative activation fold-change of EIF2AK2 in HEK293 producing PLG compared to all other transgenes in the panel. As EIF2AK2 encodes a protein involved in ER stress and the UPR, causing inhibition of mRNA translation, this remarkably lower differential expression activation of EIF2AK2 in HEK293 cells expressing PLG, along with non-detectable secretion of the PLG r-protein, may suggest that PLG expression is limited already at steps prior to ER processing in HEK293 cells. On the other hand, as HSPA8 and ERN1 showed a more consistent correlation across transgenes these genes may be universal secretory pathway markers for secreted titer differences between the cell lines across a variety of r-proteins (**Figure 6**).

At pathway levels, all cell lines responded to transgene expression by up-regulating ribosomal activities (**Figure 5C**). However, CHO cells struggled with activating the components responsible for the downstream processing and export of the r-proteins. which may explain why overexpressing a single gene in most cases only had minor positive effects or failed to boost the expression of THBS4, as the genes in pathways showing deficient expression in CHO cells likely work in tandem, and the overexpression of just one gene is often not sufficient to activate these under-expressed pathways.

While some secretory pathway genes identified to have profound positive effects on r-protein secretion seemed to assist r-protein expression indiscriminately, mounting evidence has suggested a product-specific role for many of the components within the secretory pathway. Genetic perturbation studies targeting the secretory pathway revealed that different secreted proteins utilize distinct sets of secretory pathway components during their production (Fischer et al., 2015; Ikawa et al., 1997; Leung-Hagesteijn et al., 2013; Sheng et al., 2017). Further lending credence to this product-specific nature of the secretory pathway was the secreted protein-dependent expression of the secretory pathway components. For example, protein disulfide isomerase (PDI) expression and disulfide-rich protein secretion rates are correlated across human tissues (Feizi et al., 2017) and mouse lymphocytes (Roth and Koshland, 1981). While our results show no significant PDI-dependent titer improvements, the availability of N- and O-linked glycosyltransferases can restrict titer improvements of glycosylation-enriched r-proteins (**Figure 5D**). Notably, increased glycosyltransferase activities seemed to decrease the expression of lightly- and non-glycosylated r-proteins, suggesting that when underutilized, the metabolic costs of certain cellular resources can easily outweigh their benefits in protein secretion. This cautions against the pursuit of an omnipotent host cell line and highlights the importance of customizing engineering strategies according to the properties of the r-proteins. Other key factors that can have protein-specific impacts on the secreted protein titers are e.g. RNA instability (Graf et al., 2004; Hung et al., 2010; Scholten et al., 2006), the choice of signal peptide within the transgene sequence (Dalton and Barton, 2014; Güler-Gane et al., 2016) and proneness to proteolytic degradation (Dorai et al., 2011; Gao et al., 2011; Goldman et al., 1997). Such issues should be addressed for each specific r-protein for thorough bioproduction optimization.

In summary, we show that HEK293 can serve as a valuable fall-back expression strategy, for difficult or non-secreting proteins expressed in CHO cells, or that comparisons between the different host cells can guide efforts to rescue poor expression in CHO. Taken together the results of this study shine light on the variation in expression and activation of secretory pathway related genes between HEK293 and CHO. Such cell line-specific variations could have an impact on the optimal choice of bioproduction host for specific r-proteins depending on the requirement for specific secretory pathway processing. We hypothesize that the existence of a collection of secretory machinery that better conforms to our panel of human proteins in the HEK293 cell lines is key to their improvements in protein titers. Indeed, amongst the most profound differences in expression between HEK293 and CHO secretory pathways, genes with especially positive impact on protein secretion in CHO were found. Although many of the secretory machinery components promiscuously assist the secretion of different proteins (Saibil, 2013), there are reports of more product-specific improvements to the secretory machinery (Butz et al., 2003; Ikawa et al., 1997; Yoshida et al., 2005). Supporting this, results highlighted the N- and O-linked glycosyltransferases as a group of genes aiding, or restricting, protein secretion in a protein specific manner. These highly product-specific genes enable bespoke cell line designs that cater to the unique secretory requirements of different r-proteins, and allows for a more rational selection of cell hosts for a given r-protein.

## Materials and Methods

### Experimental Design

This study was performed in three main steps. Initially, difficult to express r-proteins were produced in various expression systems (various cell lines and protocols) to evaluate performance differences in protein-specific or unspecific secreted production as determined by absorbance at 280 nm of purified r-proteins or western blotting and LC-MS/MS analysis of cell culture supernatants. In a second step, a transcriptomic evaluation of cell lines with distinct differences in performance was conducted in order to evaluate underlying secretory pathway components with impact on r-protein secretion, including differential expression analysis, gene set enrichment analysis and protein feature analysis. Genes of interest, identified by the transcriptomic profiling were subjected to co-expression analysis together with a difficult to express protein in CHO and HEK293 to evaluate the impact on r-protein secretion. ELISA or western blotting was used to determine relative r-protein titers compared to expression of the difficult to express protein alone.

### Cell lines and medium

ExpiCHO-S (Gibco™) cells were cultivated in ExpiCHO expression medium. 293-F (Gibco™) and Freestyle™ 293-F (Gibco™) cells were cultivated in FreeStyle™ 293 expression medium (Gibco™). FreeStyle™ CHO expression medium (Gibco™) supplemented with 8 mM GlutaMAX™ (Gibco™) was used for Freestyle™ CHO-S cells (Gibco™). Cells were cultivated in 125 ml Erlenmeyer shake flasks at 37°C, 8°C CO_2_ and 125 rpm. CHOEBNALT85, CHOEBNALT85-1E9 and 293ALL cells were cultivated according to manufacturer’s recommendations (Icosagen Cell Factory OÜ, Tartu, Estonia).

### Plasmids and expression constructs

For expression validation of difficult to express proteins the pQMCF vector or the in house designed pKTH16 or pKTH16_dPur plasmid (Supplemental Figure S1) was used. Expression in both vectors are driven by the CMV promoter. The pQMCF generic expression cassette included an N-terminal CD33 leader sequence (Chapple et al., 2006) followed by a short spacer sequence (AAA) and a C-terminal TEV and human protein C tag (Volk et al., 2016). The pQMCF-plasmid without gene insert served as empty vector control. Moreover the in house pKTH16 plasmid was used as expression vector with the transgene expressed fused to a N-terminal CD33 signal peptide (no spacer sequence between signal peptide and mature sequence) and a C-terminal human protein C-tag. Transgenes for co-expression validation in combination with a difficult to express protein were cloned into the pKTH16 vector in fusion with a C-terminal FLAG tag. An empty pKTH16 vector, not encoding a transgene, was used as negative control in co-expression experiments.

### ExpiCHO transfection, cultivation and harvest

The pKTH16_dPur plasmid was used for expression of transgenes. The transfections and cultivations were performed according to the manufacturers ExpiCHO standard protocol. One day prior to the transfection, the cells were seeded at 3×10^6^ cells/ml. At the day of transfection, the cells were split to 6*10^6^ cells/ml in 25 ml ExpiCHO expression medium. The ExpiFectamine reagent and 20 µg of plasmid DNA were diluted separately in OptiPRO SFM and then mixed together and incubated at room temperature for three minutes before addition to cells. The cells were cultivated at 37°C, 8% CO_2_ and 125 rpm. The day after the transfection (18-22 hours post-transfection), ExpiFectamine CHO Enhancer and ExpiCHO Feed were added to the cells. The cells were harvested at day 8 post-transfection.

### Affinity protein purification

Recombinant proteins were purified using the Anti-Protein C Affinity Matrix (11815024001, Roche) on an ASPEC liquid handling instrument (Gilson). The matrix was washed three times with equilibration buffer (20 mM Tris, 0,1 M NaCl, 2 mM CaCl_2_, adjusted to pH 7,5). The protein sample was filtered through a 0,45 µm filter and then incubated with the purification matrix overnight on a rock n roll at 4°C prior to packing of the matrix-protein mixture on columns. The column was equilibrated with 20 mM Tris, 0,1 M NaCl, 2 mM CaCl_2_, at pH 7,5 and washing was performed with 20 mM Tris, 1 M NaCl, 2mM CaCl_2_, at pH 7,5. HPC4-tagged proteins were eluted by EDTA (20 mM Tris, 0,1 M NaCl, 5 mM EDTA, pH 7,5. The elution fractions were loaded on a SDS-PAGE gel to examine the purity and the yield of the elution fractions. The elution fractions showing strong and pure bands were desalted and buffer exchanged to 1xPBS. Protein concentrations were determined by absorbance measurements at 280 nm.

### Medium-scale episomal stable expression (pQMCF system), cultivation and harvest in CHO and HEK293

293ALL and CHOEBNALT85-1E9 cells 007 (Icosagen, Tartu, Estonia) were transfected using Reagent 007 (Icosagen, Tartu, Estonia) and cultivations were performed at 30-35 ml scale at 37°C for the first three days, followed by incubation at 30°C until end of cultivation. Supernatant and cells were harvested at day 7 for 293ALL and day 9 for CHOEBNALT85-1E9. The supernatant was clarified by centrifugation at 1800 *xg* for 45 min at 20°C. Cells were stored in RNAlater™ stabilization solution (Invitrogen™) for downstream RNA extraction.

### Small-scale transient transfection, cultivation and harvest in CHO and HEK293

The pQMCF plasmids encoding 22 difficult to express target proteins and the pKTH16 plasmid encoding NRTN and NRTN pp were used for transient expression in Freestyle™ CHO, 293-F™ and Freestyle™ 293-F. At 24 hours before transfection, cells were split to 0.6 (Freestyle™ CHO) or 0.7 (293-F and Freestyle™ 293-F) million cells/ml in 125-ml Erlenmeyer shake flasks (Corning). On the day of transfection, the culture medium was exchanged by centrifugation and cells were resuspended in fresh medium at a cell density of 1 million cells/ml. Cells were transfected with 25 kDa linear PEI (Polysciences Inc.) at a DNA:PEI ratio of 1:4 (Freestyle™ CHO) or 1:3 (293-F and Freestyle™ 293-F) where 1 µg DNA was added per 1 million cells. Each plasmid was transfected in duplicate wells. The pD2529-CMVM-03 plasmid (Atum) expressing DasherGFP was used to monitor the transfection. Cultivation was performed in 24 deep-well plates with 2 ml cell suspension per well at 37°C, 8% CO_2_ and 250 rpm in humidified incubators. At 24 h post-transfection, the transfection efficiency was monitored by flow cytometry (Gallios Flow cytometer, Beckman Coulter) of GFP-expressing cells based on mean fluorescence intensities in the FL-1 channel. At 72 hours post transfection the cell culture was harvested. Cells and supernatant from each well were separated by centrifugation (500 x *g*, 3 min). Half of the cell pellet was resuspended and stored in RNAlater™ stabilization solution (Invitrogen™) according to manufacturer’s recommendations for subsequent RNA extraction.

### Expression level evaluation and protein characterization by western blot

For western blot analysis of samples, cell pellets were initially lysed in M-PER solution (Thermo Fisher Scientific) and samples were separated by SDS-PAGE (Criterion TGX Precast gels, 4-12%, Bio-Rad) under denaturing conditions. Proteins were transferred onto PVDF membranes (Trans-Blot Turbo Transfer Pack, Bio-Rad) using the Trans-Blot Turbo Blotting System (Bio-Rad), followed by blocking of membranes with 5% milk in TBST (0.05% Tween-20). Washing of membranes was performed with TBST and r-proteins were stained using a primary anti-HPC4-antibody (0.2 µg/ml, Icosagen) followed by the secondary goat anti-human HRP-conjugated antibody (1:4000, A18805, Invitrogen). Stained proteins were detected using Immobilon Western Chemiluminescent HRP Substrate (Millipore) and image acquisition using a ChemiDoc Imaging system (Bio-rad). For protein abundance estimations, each supernatant sample was run in duplicate and volumetric band intensities were fitted onto a standard curve generated by a dilution series of HPC4-tagged EPO on the same membrane using the Image Lab software (Bio-Rad).

### Transcriptome profiling

RNA was extracted using RNeasy plus Mini Kit (Qiagen) according to manufacturer’s guidelines and the quality of the isolated RNA was evaluated by BIOanalyzer 2100 (Agilent, Santa Clara, CA) using the Agilent RNA 6000 Nano Kit. RNA sequencing was performed at GATC Biotech (Konstanz, Germany) using the Inview transcriptomics Discover platform. Sequence data for RNA-Seq were quality controlled using FastQC and summarized with multiQC (Ewels et al., 2016). Trimmomatic (Bolger et al., 2014) was used to trim low-quality bases from the reads. The CHO-K1 (Lewis et al., 2013; Rupp et al., 2018) and the human GRCh38.p12 reference genomes were extended to incorporate the transgene sequences so that the transcripts of the heterologous secretome can be quantified. Reads were then quasi-mapped to either the extended CHO-K1 or the human GRCh38.p12 genome based on the cell line of origin and quantified with Salmon (Patro et al., 2017). To compare the transcriptome usage across CHO and HEK293 cell lines, an ortholog conversion table (Kallehauge et al., 2017) was used to convert CHO genes to their human orthologs. The functional groups and the color scheme for the transcriptome usage were adapted from the Proteomaps tool (Liebermeister et al., 2014).

Differential expression was performed using DESeq2 (Love et al., 2014). To facilitate the comparison between CHO and HEK293 cells, CHO genes were converted to human orthologs using the conversion table described above. With tximport (Soneson et al., 2016), the transcript-level abundances were integrated into gene-level counts to be compatible with DESeq2. Three different types of comparisons were carried out by specifying the corresponding design matrices. For cell line comparisons, gene expression profiles from all producers were compared between HEK293 and CHO cells. To estimate the degree of gene activation upon recombinant production, the CHO and HEK293 producer cells were compared with their non-producing counterparts respectively. To better account for sample variation, the fold changes were shrunken towards a beta prior to reduce effect sizes for low confidence fold change estimates and improve gene fold changes rankings (Zhu et al., 2019), eliminating the need for additional filtering. To obtain genes that show the greatest activation disparity between HEK293 and CHO cells, the absolute difference of fold changes for all genes across the two cell lines were ranked, and the top 20 (0.12%) of them were chosen for further investigation. To estimate the differential activation between cell lines across r-proteins, the DESeq2 design matrix was expanded to include an interaction term between the cell lines and the r-protein identities.

To not be overshadowed by the expression of extreme genes and to pay more equal attention to all the genes within the secretory pathway, a variance-stabilizing transformation was applied (Anders and Huber, 2010), similar in implementation to a log-transformation on the secretory pathway gene expression.

### Co-expression validation of gene outliers between HEK293 and CHO

Each pKTH16_dPur plasmid encoding gene outliers between HEK293 and CHO, or an empty pKTH16_dPur vector control, was co-transfected with the pQMCF plasmid encoding THBS4 in duplicates into 293-F and Freestyle CHO-S cells using PEI as described above. In total 1 ug of plasmid was transfected per 1 million cells and plasmid ratios of 1:1, 1:2 and/or 1:10 (gene outlier:THBS4) was used. Cells were cultivated in deep-well plates as described above and cells and supernatant were harvested at 72 hours post transfection. Some gene outliers were also validated in combination with ARTN. The expression of THBS4 in each sample was evaluated by sandwich ELISA using a human anti-HPC4-antibody (Icosagen) as capture antibody, rabbit anti-THBS4 (antibody HPRK2400008 kindly provided from the Human Protein Atlas) as primary antibody and a HRP-conjugated swine anti-rabbit antibody (p039901-2, Dako) combined with TMB substrate (Thermo Fisher Scientific) for detection. The relative secreted expression of ARTN was determined by western blotting of culture supernatants using the rabbit antibody HPRK3140989 from the Human Protein Atlas and an HRP-conjugated swine anti-rabbit antibody (p039901-2, Dako) combined with Immobilon Western Chemiluminescent HRP Substrate (Millipore). For western blot validation of outlier gene expression, cells were lysed using M-PER or Mem-PER mammalian protein extraction reagents (Thermo Fisher Scientific) and blotted onto PVDF membranes as described above. Proteins were detected using a monoclonal anti-FLAG M2 antibody (F3165, Sigma/Merck Millipore) and an HRP-conjugated polyclonal goat anti-mouse antibody (P0447, Dako).

### Pathway and protein feature analysis

Gene set enrichment analysis was used to calculate the significantly activated pathways from gene-level differential expression profiles. Canonical pathway annotation was obtained from MSigDB (Liberzon et al., 2011). Additionally, manually curated gene sets on various secretory pathway subsystems were referenced from (Feizi et al., 2017). A normalized enrichment score (NES) representing the gene-set enrichment analysis (GSEA) statistic (Subramanian et al., 2005) was calculated to quantify the overall direction of regulation for each gene set along with an accompanying permutation p-value (Korotkevich et al., 2021).

PTM information for each r-protein integrated from UniProt and phosphosite. Among the various types of common PTMs that occur in the cell, we considered glycosylation and disulfide bonds due to their occurrences in our panel of r-proteins and their relevance to the secretory pathway. For each r-protein, a glycosylation- and disulfide bond-index quantifying the level of enrichment of the respective PTM was calculated by dividing the total number of occurrences of the PTM in question in the r-protein by the length of the protein. The enzymes responsible for the synthesis of glycans and disulfide bonds were obtained from (Narimatsu et al., 2019) and KEGG (Kanehisa and Goto, 2000) respectively, and their corresponding expression changes from CHO to HEK293 were summarized using the GSEA enrichment score statistic for each r-protein. A Bayesian linear regression model (formula given below) was used to assess the relationship between titer improvement from CHO to HEK293 and the enzyme expression. To further deconvolute the effects of PTMs on this dependency, an interaction term between the PTM index and the enzyme expression was added to the linear model to capture how the correlation between the enzyme activities and the titer improvements changes across r-proteins with different PTM indices. The model coefficients were estimated with Markov chain Monte Carlo (MCMC) via the rethinking package with default parameters (McElreath, 2020).

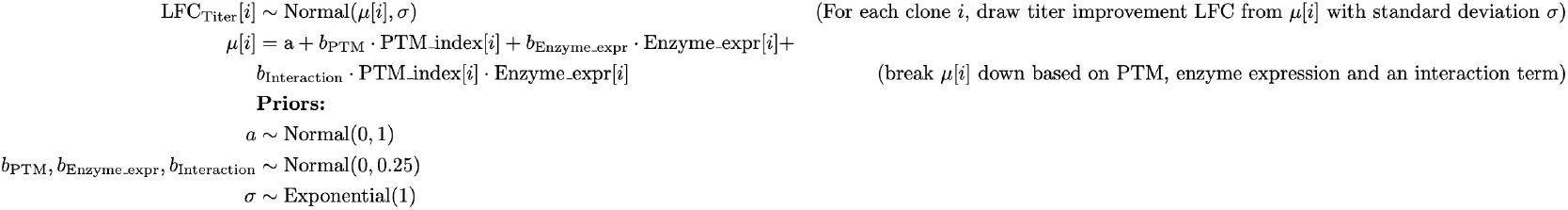

### Statistical Analysis

Statistical analysis of significantly different expression levels of THBS4 and ARTN from co-expression experiments was performed in GraphPad Prism 7 using ANOVA one-way analysis followed by Durnetts’s test comparing all expression levels to a control (THBS4 or ARTN co-expressed with an empty plasmid). The details of the statistical analysis of the transcriptomic profiling, including differential expression analysis and protein feature analysis, is described in the respective sections above.

## Author contributions

Conceptualization, Ma.M., C.K., J.R., D.H., R.F., P.V., N.E.L. and V.C.; Methodology, Ma.M., C.K., N.E.L and J.R., F.E.; Investigation, Ma.M., C.K., Mo.M., A.M., N.W., R.R., A.V., M.L., D.K, N.E.L. and J.R.; Writing – Original Draft, Ma.M, C.K., Mo.M. and J.R.; Writing – Review & Editing, Ma.M., C.K., T.G., D.H., N.E.L. and J.R.; Visualization, Ma.M., C.K. and Mo.M; Supervision, J.R., N.E.L., D.H., and P.V.; Funding Acquisition, V.C., R.F., P.V., N.E.L. and J.R.

## Data and materials availability

Raw RNA sequencing data are deposited to the Sequence Read Archive (SRA) with accession number SRP281874, and the corresponding processed data are available on the Gene Expression Omnibus (GEO) via accession number GSE157729. All the other data are contained in the article and its supplementary information or available upon request. The source code for the figures is available from https://github.com/LewisLabUCSD/CHO_HEK.

