## Supplemental data for "Harnessing secretory pathway differences between HEK293 and CHO to rescue production of difficult to express proteins"

**Supplemental Text**

***LC-MS/MS analysis using SIS PrESTs and QTag standards***

The LC-MS/MS standards were produced as SIS PrESTs (Edfors et al., 2014)., produced in auxotrophic E. coli strain (Zeiler et al., 2012), were purified by the standard workflow used within the Human Protein Atlas for PrEST production (Tegel et al., 2009) and quantified by bicinchoninic acid assay. Every SIS PrEST has a purification tag HisABPOneStrep (QTag), that can also be used for their quantification by mass spectrometry using its light version. The light QTag was quantified by amino acid analysis. Therefore, the target protein can be precisely quantified in a sandwich manner where SIS PrEST is quantified by QTag and a target protein by the overlapping peptides of SIS PrEST in a single MS run (Fig. A).

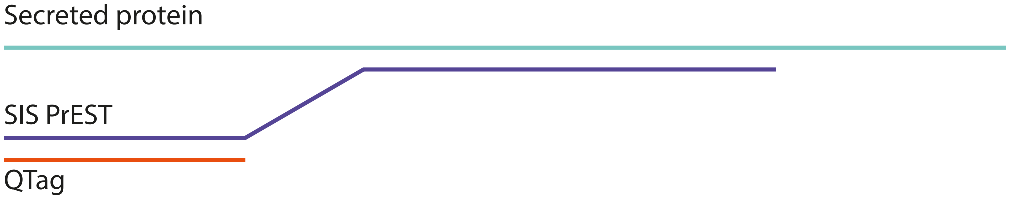


Figure A: Strategy for a sandwich quantification of secreted proteins using amino acid-analyzed QTag.

Equimolar amounts of SIS PrESTs were pooled (Table S-2), reduced (2mM DTT, 30 min, 56 °C), alkylated (10mM CAA, 20 min in the dark at room temperature (RT)) and digested with proteomics grade porcine trypsin (Sigma Aldrich) in a 1:50 (w/w) ratio of enzyme to substrate (37 °C, 300 rpm, O/N). Enzymatic reaction was quenched with 0.5% (v/v) formic acid and samples vacuum dried and stored at -20 °C.

Initially, 100 µl of the cell supernatants (~30ug of protein content, determined by Bradford protein assay (Bio-Rad)) were aliquoted and spiked-in with equimolar mixture of corresponding SIS PrEST and QTag (Table S-2). Protein mixture was precipitated with four times the volume of ice-cold acetone (O/N, -20 °C). Precipitate was centrifuged (30 min, 4 °C, 20 000 rcf), supernatant discarded and pellet washed with 400 µl of ice-cold acetone followed by centrifugation (10 min, 4 °C, 20 000 rcf). Pellet was dried and dissolved in 20 µl of 7M Urea, 2M Thiourea. After 8x dilution with 100mM TEAB, proteins were reduced (10mM DTT, 1h at 30 °C), alkylated (50mM CAA, 30 min, RT in the dark) and digested with 600 ng of porcine trypsin (Sigma-Aldrich) O/N at 37 °C. Enzymatic reaction was quenched with 0.5% (v/v) trifluoroacetic acid, samples solid-phase extracted using in-house prepared C18 StageTips (Edfors et al., 2018), vacuum dried and stored at -20 °C.

Samples were analyzed using the UltiMate 3000 nano-liquid chromatography system (Thermo Scientific) with an EASY-Spray ion source connected to Q Exactive HF (Thermo Scientific) mass spectrometer. Samples were resuspended in solvent A (3% ACN, 97% H2O, 0.1% FA) and peptides loaded onto an Acclaim PepMap 100 trap column (75 µm × 2 cm, C18, 3 µm, 100 Å, Thermo Scientific), washed 5 min at 5 µl/min with 100% of solvent A and separated with

PepMap RSLC C18 (75 µm x 25 cm, 2 µm, 100 Å, Thermo Scientific) analytical column using linear 45 min gradient of 6-37% solvent B (95% ACN, 5% H20, 0.1% FA) at a flow rate of 0.400 µl/min and analyzed in either Top10 or parallel reaction monitoring (PRM) mode. The analytical and trap columns were kept at 35 °C by the in-source temperature controller and 40 °C by the column oven temperature controller respectively.

Samples were resuspended in solvent A and peptides corresponding to 500 fmol/SIS PrEST were injected and analyzed using a Top10 MS-method. MS1 scan, performed at resolution of 60,000 (mass range 400–1,400 m/z, AGC 3e6), was followed by ten consecutive MS2 scans at resolution of 30,000 (AGC 1e6, Max IT 150 ms) with normalized collision energy set to 27. Resulting raw files were searched using MaxQuant (version 1.5.3.30) (Cox and Mann, 2008) with the built in search engine Andromeda against whole human reference proteome downloaded from Uniprot (Swissprot, 18th September 2017, UP000005640, 20,205 entries). Search parameters were set to peptide length ranging 7-25 amino acids, enzyme specificity: trypsin with maximum of two miscleavages, 4.5 ppm match tolerance for precursor ions and 20 ppm for fragment ions with 1% false discovery rate on both the peptide and protein level. Maximum of five variable modifications by oxidation on methionine and N-terminal acetylation were allowed and carbamidomethylation on cysteine was set as a fixed modification. Arg10 and Lys8 were chosen as heavy labels with a maximum of 3 labels per peptide. Resulting msms files were used to build a spectral library in open source targeted proteomics environment Skyline (Maclean et al., 2010) .

Supernatant samples were resuspended by the autosampler and peptides corresponding to 2 µg of protein content injected and analyzed in parallel reaction monitoring (PRM) mode with inclusion list containing either all theoretical SIS PrEST peptides with no methionine and the samples were later rerun with only detected peptides or all peptides that were found using the SIS PrEST library in Skyline. All inclusion lists also contained QTag peptide ISEATDGLSDFLK that was used for a precise SIS PrEST quantification (Table S-3). The method for PRM mode consisted of one full MS scan at resolution of 60,000 (AGC target 3e6, mass range 200-1,600 m/z and injection time 110 ms) followed by 20 MS2 scans at resolution of 60,000 (AGC target 2e5, isolation window 1.2 m/z, injection time 110 ms and NCE 27). Resulting raw files were imported to Skyline (Maclean et al., 2010) and peptide ratios extracted and statistically analyzed.

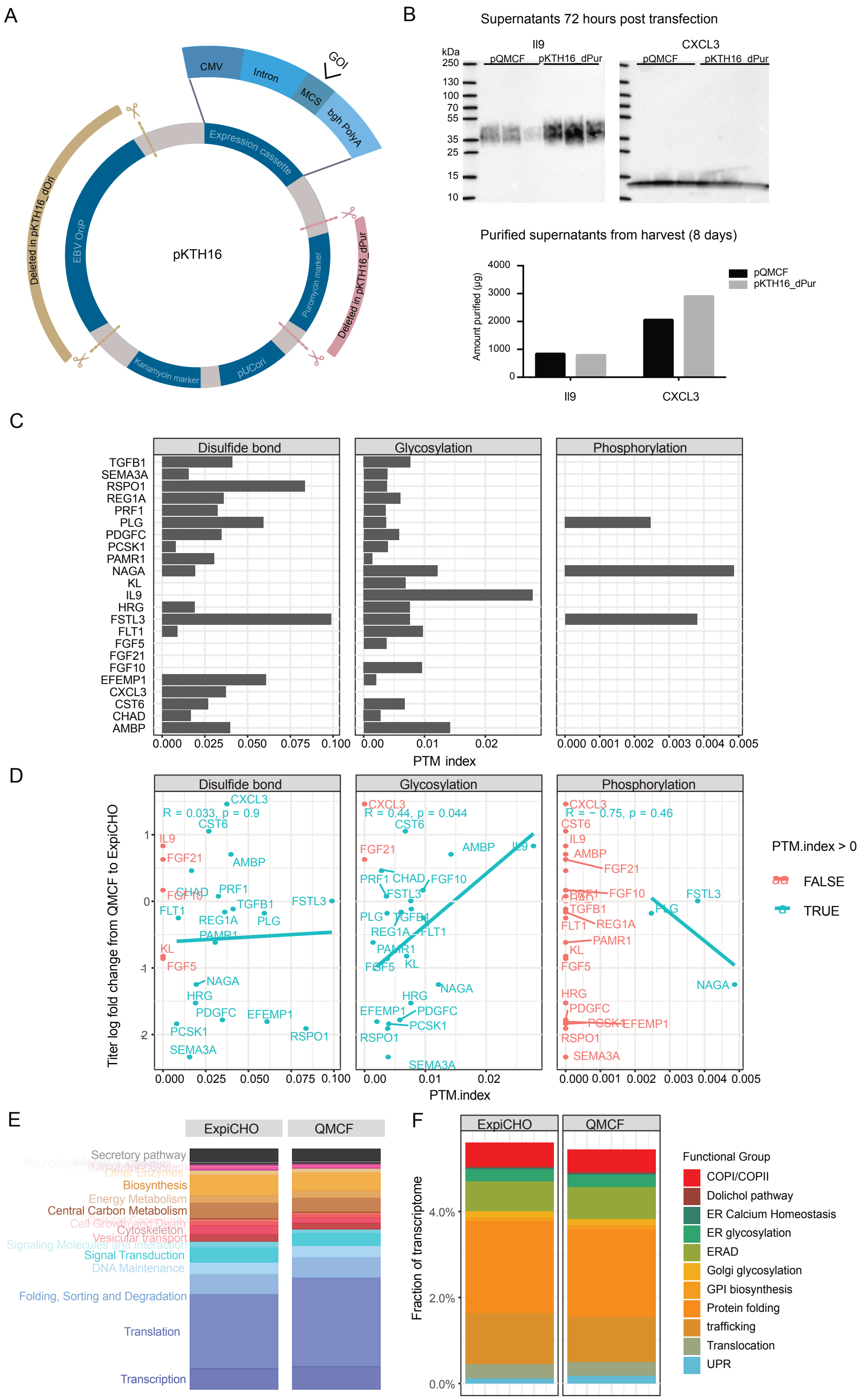

A
CMV
Intron
GOI
MCS
bgh PolyA
Expression cassette
Deleted in pKTH16_dOri
EBV OriP
pKTH16
Puromycin marker
Deleted in pKTH16_dPur
Kanamycin marker
pUCori
B
Supernatants 72 hours post transfection
Il9
CXCL3
kDa
pQMCF
pKTH16_dPur
250
130
100
70
55
35
25
15
10
Purified supernatants from harvest (8 days)
Amount purified (µg)
4000
3000
2000
1000
0
Il9
CXCL3
C
Disulfide bond
Glycosylation
Phosphorylation
TGFB1
SEMA3A
RSPO1
REG1A
PRF1
PLG
PDGFC
PCSK1
PAMR1
NAGA
KL
IL9
HRG
FSTL3
FLT1
FGF5
FGF21
FGF10
EFEMP1
CXCL3
CST6
CHAD
AMBP
0.000 0.025 0.050 0.075 0.100
0.00 0.01 0.02
0.000 0.001 0.002 0.003 0.004 0.005
PTM index
D
Titer log fold change from QMCF to ExpiCHO
R = 0.033, p = 0.9
R = 0.44, p = 0.044
R = − 0.75, p = 0.46
PTM.index
PTM.index > 0
FALSE
TRUE
E
ExpiCHO
QMCF
Secretory pathway
Other Enzymes
Biosynthesis
Energy Metabolism
Central Carbon Metabolism
Cytoskeleton
Vesicular transport
Signal Transduction
DNA Maintenance
Folding, Sorting and Degradation
Translation
Transcription
F
Fraction of transcriptome
4.0%
2.0%
0.0%
Functional Group
COPI/COPII
Dolichol pathway
ER Calcium Homeostasis
ER glycosylation
ERAD
Golgi glycosylation
GPI biosynthesis
Protein folding
trafficking
Translocation
UPR

**Supplemental Figure S1.** Related to Figure 1.

(A) An overview of the in house designed pKTH16 plasmid. The plasmid contains an expression cassette with the CMV promoter, a modified intron A from human CMV, a multiple cloning site (MCS) for the gene of interest (GOI) followed by the poly(A) signal from bovine growth hormone (bgh). The design of the plasmid allows for easy restriction enzyme-based removal of the puromycin marker gene and/or the oriP from Epstein bar virus (EBV) generating pKTH16_dPur and/or pKTH16_dOri, respectively. (B) Evaluation of expression of IL9 and CXCL3 using the pKTH16_dPur plasmid was compared to expression using the pQMCF plasmid in a fully transient mode. Western blots from culture supernatants, at 72 hours post transfection, of triplicate cultures of each protein and expression vector and the purified titer (from the replicate with the highest protein content) after 8 days of cultivation confirmed equal or better expression obtained using pKTH16 compared to pQMCF in a fully transient expression mode. (C) The level of enrichment of three common PTMs in each recombinant protein. In each panel, the length of the bar represents the PTM index for a given recombinant protein, or the total number of known PTM occurrences divided by the length of the protein. (D) Scatter plot comparing the PTM index and the titer fold changes from the QMCF to the ExpiCHO platform (positive fold changes indicate higher titers in the ExpiCHO system). Each panel shows the correlation for one of the PTMs, where correlation coefficient is calculated based on clones harboring the given PTM (non-zero PTM index). (E) Comparison of overall transcriptome usage between the ExpiCHO and the QMCF expression systems. The fraction of the transcriptome dedicated to various cellular functions is represented by the height of each bar, where related pathways are colored similarly. (F) Transcriptome usage across the two expression systems, broken down by secretory pathway sub-groups.

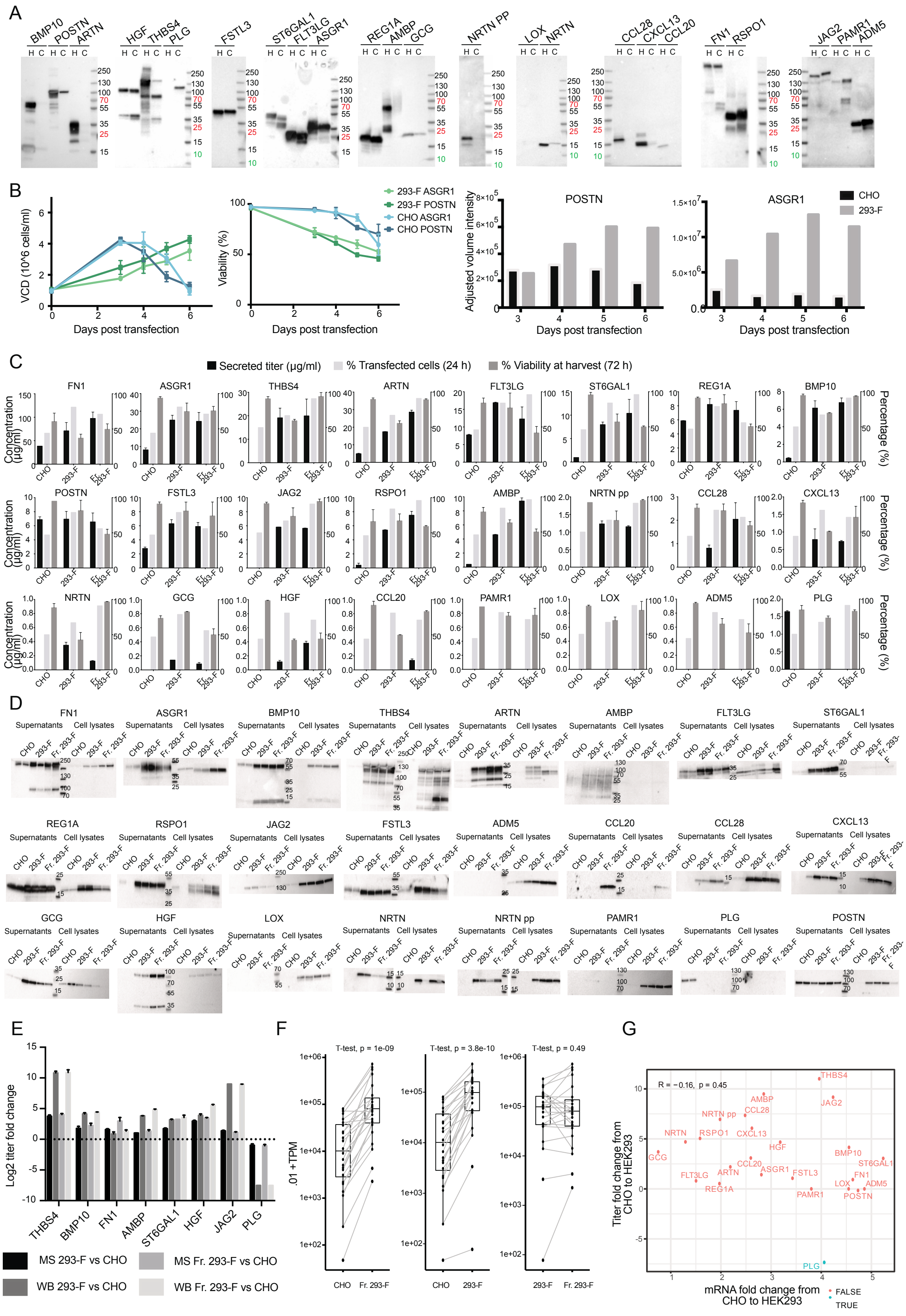
A
BMP10 POSTN ARTN
HGF THBS4 PLG
FSTL3
ST6GAL1 FLT3LG ASGR1
REG1A AMBP GCG
NRTN PP
LOX NRTN
CCL28 CXCL13 CCL20
FN1 RSPO1
JAG2 PAMR1 ADM5
H C
250
130
100
70
55
35
25
15
10
B
VCD (10^6 cells/ml)
Days post transfection
Viability (%)
293-F ASGR1
293-F POSTN
CHO ASGR1
CHO POSTN
Adjusted volume intensity
POSTN
ASGR1
CHO
293-F
C
Secreted titer (µg/ml)
% Transfected cells (24 h)
% Viability at harvest (72 h)
Concentration (µg/ml)
Percentage (%)
CHO
293-F
Fr. 293-F
FN1 ASGR1 THBS4 ARTN FLT3LG ST6GAL1 REG1A BMP10
POSTN FSTL3 JAG2 RSPO1 AMBP NRTN pp CCL28 CXCL13
NRTN GCG HGF CCL20 PAMR1 LOX ADM5 PLG
D
Supernatants
Cell lysates
FN1 ASGR1 BMP10 THBS4 ARTN AMBP FLT3LG ST6GAL1
REG1A RSPO1 JAG2 FSTL3 ADM5 CCL20 CCL28 CXCL13
GCG HGF LOX NRTN NRTN pp PAMR1 PLG POSTN
E
Log2 titer fold change
THBS4 BMP10 FN1 AMBP ST6GAL1 HGF JAG2 PLG
MS 293-F vs CHO
MS Fr. 293-F vs CHO
WB 293-F vs CHO
WB Fr. 293-F vs CHO
F
T-test, p = 1e-09
T-test, p = 3.8e-10
T-test, p = 0.49
.01 + TPM
CHO
Fr. 293-F
CHO
293-F
293-F
Fr. 293-F
G
R = −0.16, p = 0.45
Titer fold change from CHO to HEK293
mRNA fold change from CHO to HEK293
FALSE
TRUE
THBS4 JAG2 AMBP CCL28 NRTN pp NRTN RSPO1 CXCL13 GCG HGF BMP10 ST6GAL1 CCL20 ASGR1 FSTL3 FN1 ARTN FLT3LG LOX ADM5 REG1A PAMR1 POSTN PLG

**Supplemental Figure S2.** Related to Figure 2.

Expression of difficult to express proteins in HEK293 and CHO, related to Figure 2. (A) Western blots of supernatants from episomal stable expression cultures (C = CHOEBNALT85-1E9; H = 293ALL). (B) Transient expression protocol setup. Viable cell density (VCD), viability (mean values of biological replicates ± SD) and r-protein expression of Freestyle CHO-S and 293-F cells expressing ASRG1 or POSTN during cultivation for up to six days**.** Volumetric intensities of stained protein (POSTN or ASGR1) were determined by western blotting of samples harvested each day post transfection. (C) Mean values of biological replicates ± SD on secreted protein concentrations (in supernatants harvested 72 hours post transfection), transfectivity of cells (as determined by percentage of GFP positive cells in a control sample at 24 hours post transfection) and percentage of viable cells at harvest (72 hours post transfection) for each of the 24 transiently expressed transgenes in Freestyle CHO-S (CHO), 293-F and Freestyle 293-F (Fr. 293-F). (D) Western blots of supernatants and cell lysates from transient HEK293 (293-F and Freestyle 293-F) and CHO (Freestyle CHO-S) cultures expressing difficult to express proteins. (E) Titer fold changes between HEK293 and CHO determined by either MS/MS and SIS prEST technology (mean values of technical replicates ± SD) or western blot (WB) (mean values of biological replicates ± SD) for a subset of supernatant samples of the three cell lines Freestyle CHO-S, 293-F and Freestyle 293-F (Fr. 293-F). (F) Comparison of transgene mRNA abundance across several CHO and HEK cell lines. Both 293-F and Freestyle 293-F (Fr. 293-F) cells saw a significant increase in transgene transcript abundance. (G) Protein titer and mRNA changes from CHO to HEK293. No significant correlation was seen between the mRNA and titer fold changes from CHO to HEK293.

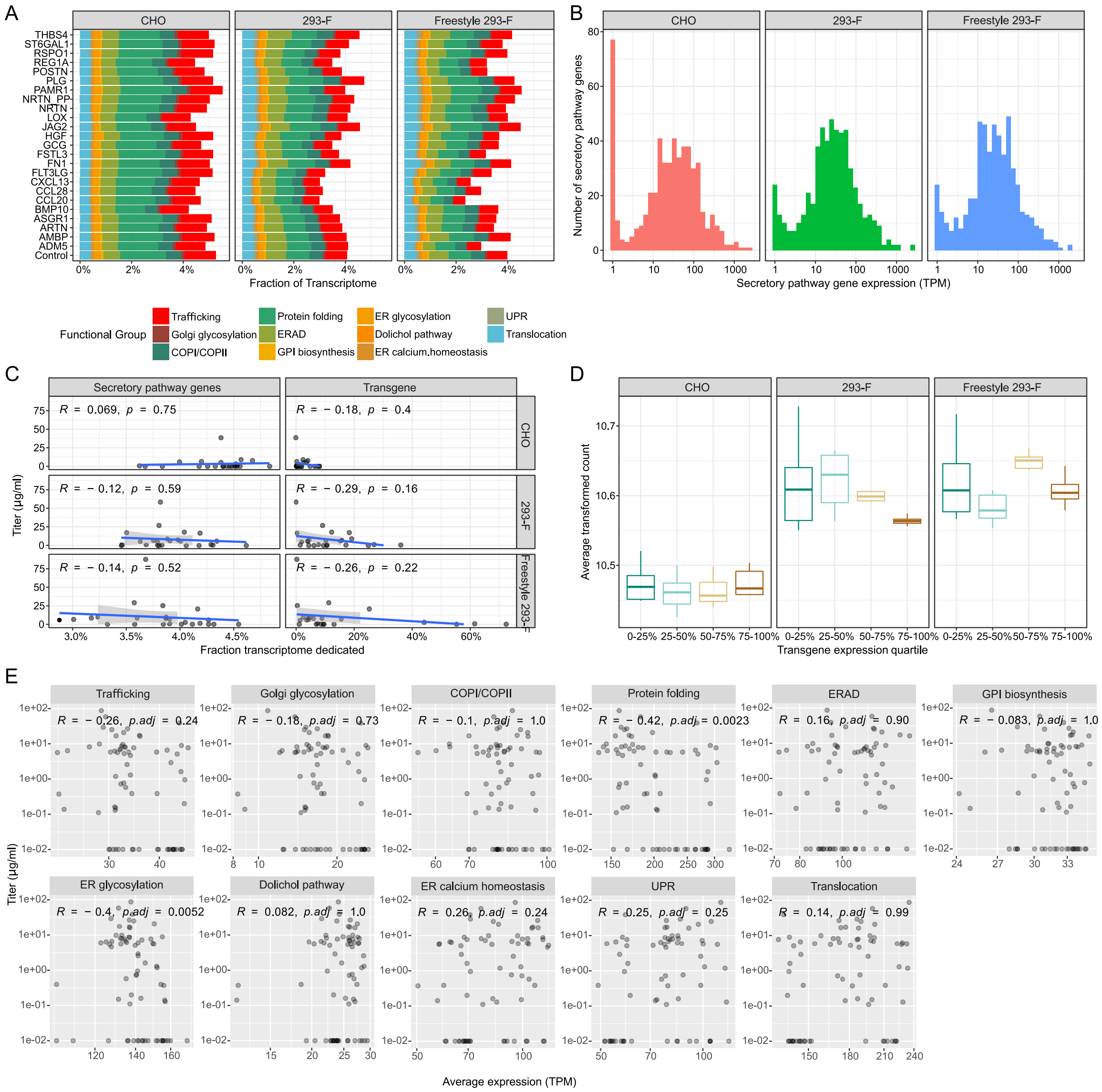

**Supplemental Figure S3.** Related to Figure 3.
(A) Transcriptome usage, broken down by secretory pathway sub-groups. Overall, the CHO cells show elevated secretory pathway activities compared to the two variants of the HEK cell lines. (B) Histogram of secretory pathway gene expression (TPM) across cell lines. Note that CHO showed a more left-skewed distribution compared to the two HEK293 variants as a result of having more non-expressed secretory pathway components, with 10.3% of the secretory pathway genes in CHO non-expressed compared to 1.9% for HEK293 cells. (C) Correlation between r-protein titers (y-axis) and fraction of transcriptome dedicated (x-axis) to secretory pathway genes (left panels) and transgene (right panels) in each cell line. (D) Average secretory pathway gene expression (y-axis, in transformed counts) across clones with different transgene loads (x-axis), where the producers were divided into 4 quartiles according to their transgene expression levels. A linear relationship between expression of several secretory pathway components and transgene mRNA abundances can be seen in CHO producers but not in HEK293 cells, where peak

secretory pathway activities occur in clones with low to medium transgene load**.** This suggests a saturation of the secretory pathway in HEK293 cells, which may be a result of the exceptionally high transgene mRNA loads observed in case of several transgenes. (E) Scatter plot for r-protein titer (y-axis) vs. average gene expression (x-axis) for each functional group within the secretory pathway. Spearman correlation coefficient and corresponding FDR adjusted p-value are given for each functional group. Only groups surviving multiple hypothesis testing are protein folding and ER glycosylation.

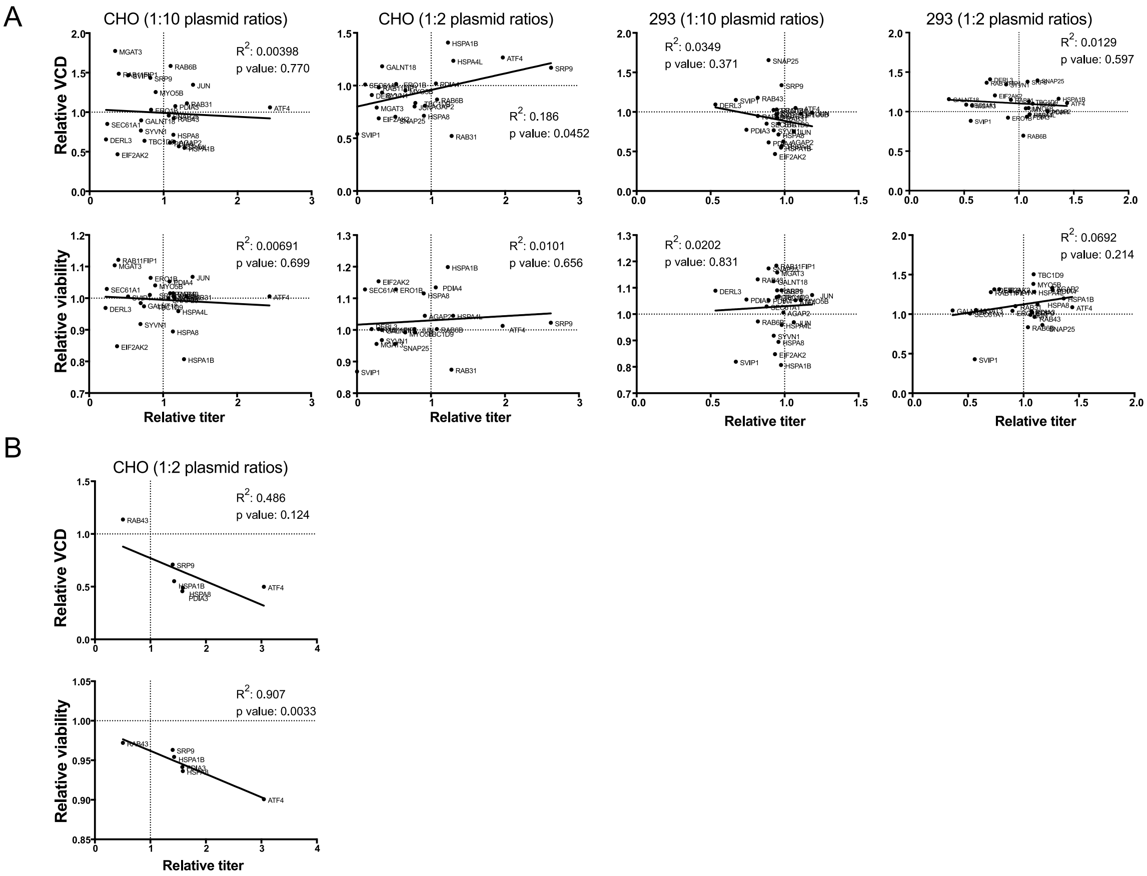


**Supplemental Figure S4.** Related to Figure 4.

Viable cell density (VCD) or viability compared to titer of (A) THBS4 and (B) ARTN at harvest of co-expression samples, related to Figure 4. Values are given as relative levels for each secretory pathway gene (labels in plot) co-expressed with THBS4 compared to controls (THBS4 co-expressed with empty vector).

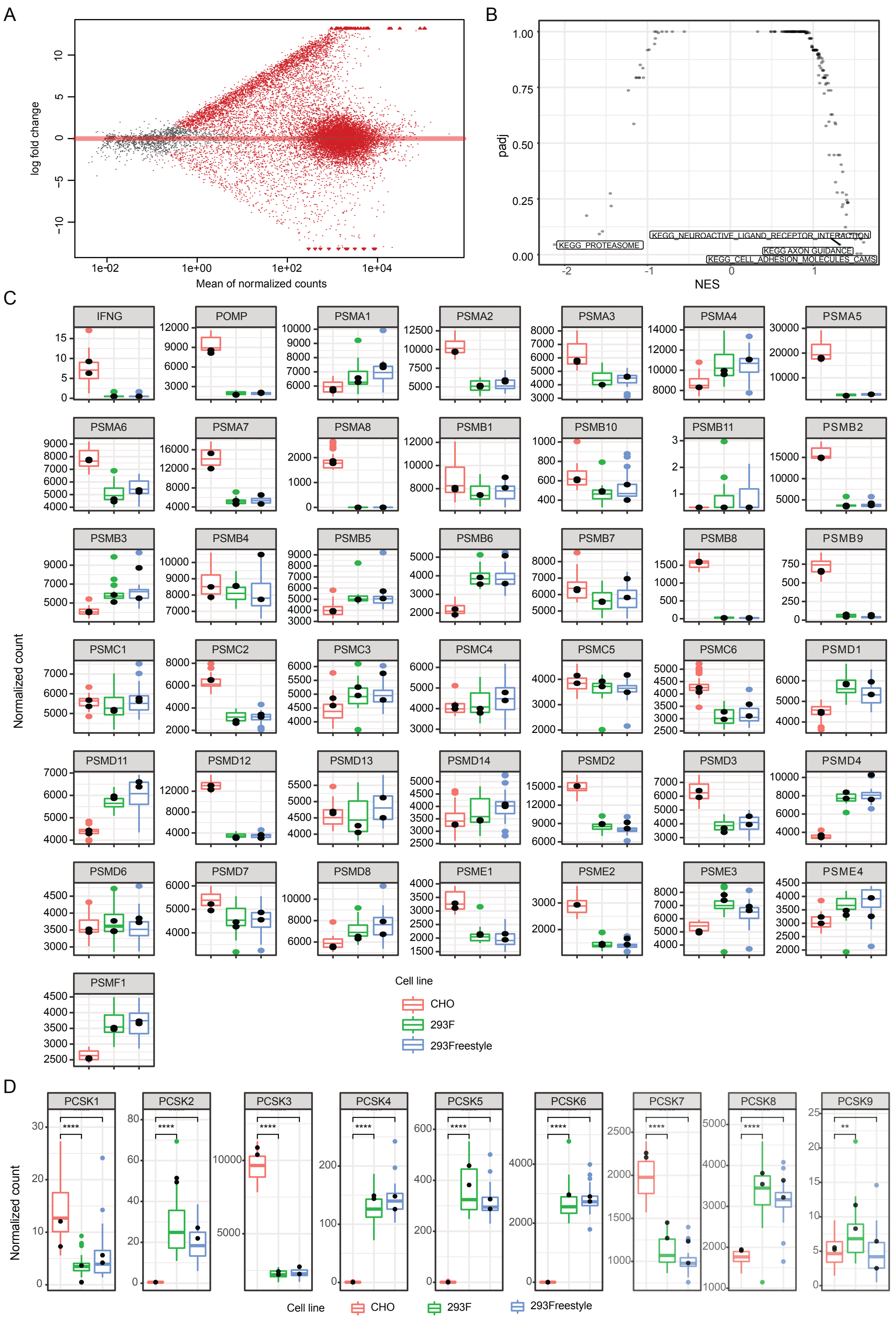
A
log fold change
Mean of normalized counts
B
padj
NES
KEGG_PROTEASOME
KEGG_NEUROACTIVE_LIGAND_RECEPTOR_INTERACTION
KEGG AXON GUIDANCE
KEGG_CELL_ADHESION_MOLECULES_CAMS
C
Normalized count
IFNG
POMP
PSMA1
PSMA2
PSMA3
PSMA4
PSMA5
PSMA6
PSMA7
PSMA8
PSMB1
PSMB10
PSMB11
PSMB2
PSMB3
PSMB4
PSMB5
PSMB6
PSMB7
PSMB8
PSMB9
PSMC1
PSMC2
PSMC3
PSMC4
PSMC5
PSMC6
PSMD1
PSMD11
PSMD12
PSMD13
PSMD14
PSMD2
PSMD3
PSMD4
PSMD6
PSMD7
PSMD8
PSME1
PSME2
PSME3
PSME4
PSMF1
Cell line
CHO
293F
293Freestyle
D
Normalized count
PCSK1
PCSK2
PCSK3
PCSK4
PCSK5
PCSK6
PCSK7
PCSK8
PCSK9
Cell line
CHO
293F
293Freestyle

**Supplemental Figure S5.** Related to Figure 5.
(A) MA plot for the differential expression between the HEK293 and CHO cell lines. A positive fold change indicates higher expression in the HEK293 cells. Red dots represent significantly differentially expressed genes (FDR corrected p-value < 0.05). (B) Gene set enrichment analysis of the differentially expressed genes between HEK293 and CHO. KEGG pathways were used, and only those that result in a FDR corrected p-value. (C) Expression comparison of 43 proteasomal genes across HEK and CHO cell lines. (D) Expression comparison of 9 common propeptide convertases across HEK and CHO cell lines.

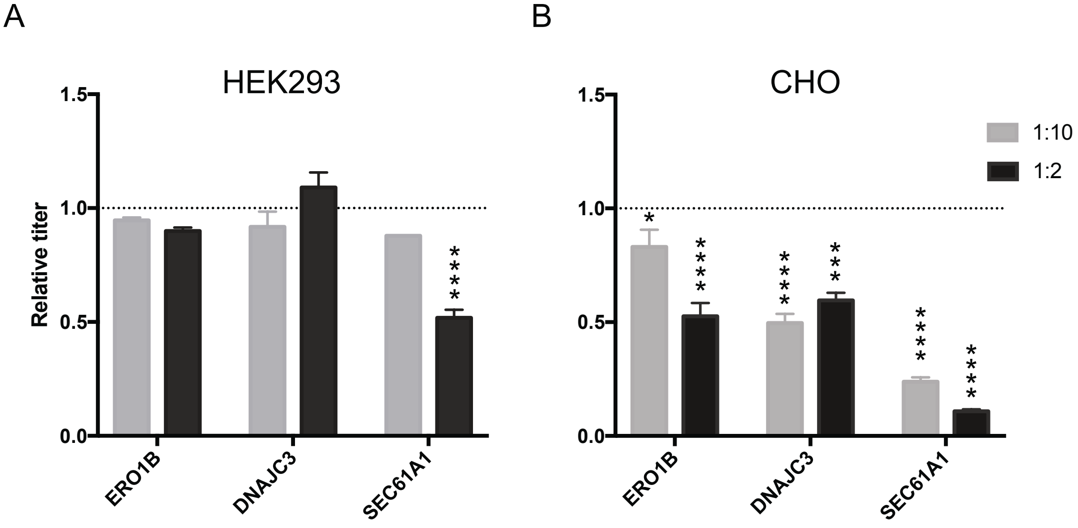


**Supplemental Figure S6.** Related to Figure 5. Effect of overexpression of differentially activated genes ERO1B, DNAJC3 and SEC61A1 on secreted production of THBS4 in A) HEK293 and B) CHO. Results are provided as relative ratios of secreted titers as compared to co-expression of THBS4 with an empty vector. Co-expression was performed with differentially activated gene to THBS4 transgene plasmid ratios of 1:2 or 1:10. Significant different expression of THBS4 (N = 2) compared to the co-expression with an empty vector control (determined by one-way ANOVA and Dunnett's test) are indicated by the asterisk sign (* P adj ≤ 0.05; ** P adj ≤ 0.01; *** P adj ≤ 0.001; **** P adj ≤ 0.0001).

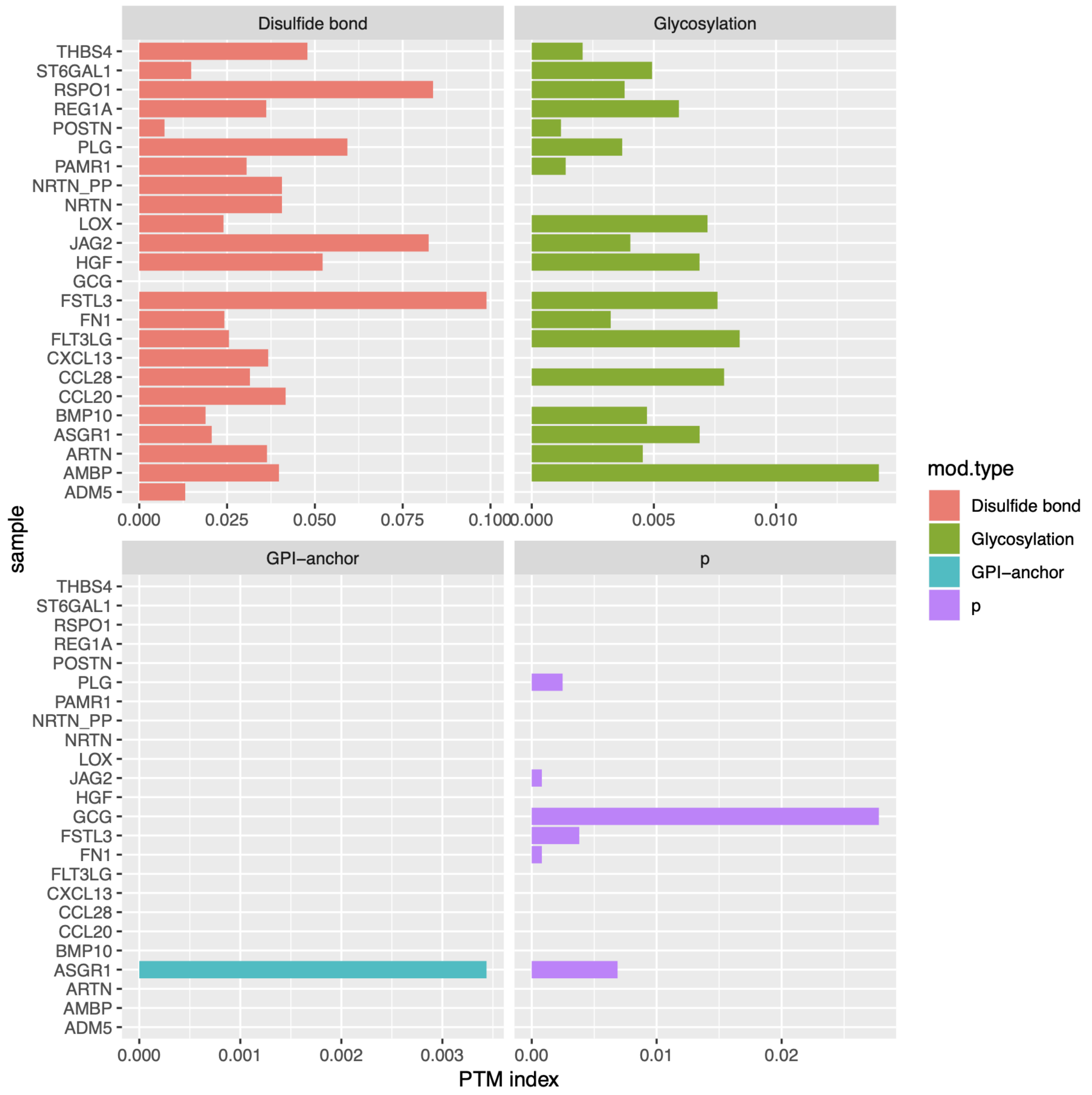


**Supplemental Figure S7.**

The level of enrichment of four common PTMs in each of recombinant protein. In each panel, the length of the bar represents the PTM index for a given recombinant protein, or the total number of known PTM occurrences divided by the length of the protein.

**Supplemental Table S1:** Amounts of spiked-in SIS PrEST and QTag standards into corresponding producers.

| SIS PrEST ID | Gene | Cell line | Standards spike-in [pmol] |
|---|---|---|---|
| HPRR2320064 | PLG | CHO | 10 |
| HPRR2320064 | PLG | 293F | 10 |
| HPRR2320064 | PLG | 293Freestyle | 10 |
| HPRR2520012 | JAG2 | CHO | 1 |
| HPRR2520012 | JAG2 | 293F | 1 |
| HPRR2520012 | JAG2 | 293Freestyle | 1 |
| HPRR2400003 | THBS4 | CHO | 0.5 |
| HPRR2400003 | THBS4 | 293F | 20 |
| HPRR2400003 | THBS4 | 293Freestyle | 20 |
| HPRR3140016 | HGF | CHO | 0.2 |
| HPRR3140016 | HGF | 293F | 2 |
| HPRR3140016 | HGF | 293Freestyle | 2 |
| HPRR1951412 | BMP10 | CHO | 1 |
| HPRR1951412 | BMP10 | 293F | 1 |
| HPRR1951412 | BMP10 | 293Freestyle | 1 |
| HPRR3720239 | ST6GAL1 | CHO | 10 |
| HPRR3720239 | ST6GAL1 | 293F | 10 |
| HPRR3720239 | ST6GAL1 | 293Freestyle | 10 |
| HPRR2050096 | FN1 | CHO | 10 |
| HPRR2050096 | FN1 | 293F | 10 |
| HPRR2050096 | FN1 | 293Freestyle | 10 |
| HPRR350094 | AMBP | CHO | 10 |
| HPRR350094 | AMBP | 293F | 10 |
| HPRR350094 | AMBP | 293Freestyle | 10 |

**Supplemental Table S2:** Precursor masses and charges of peptides used for the quantification of secreted proteins and SIS PrESTs

| Protein | Mass [m/z] | CS [z] | Comment |
|---|---|---|---|
| Qtag | 698.353739 | 2 | ISEATDGLSDFLK (light) |
| Qtag | 702.360838 | 2 | ISEATDGLSDFLK (heavy) |
| AMBP | 745.648091 | 3 | KEDSC[+57.021464]QLGYSAGPC[+57.021464]MGMTSR (light) |
| AMBP | 751.65558 | 3 | KEDSC[+57.021464]QLGYSAGPC[+57.021464]MGMTSR (heavy) |
| AMBP | 607.339715 | 2 | TVAAC[+57.021464]NLPIVR (light) |
| AMBP | 612.343849 | 2 | TVAAC[+57.021464]NLPIVR (heavy) |
| AMBP | 835.874324 | 2 | C[+57.021464]VLFPYGGC[+57.021464]QGNGNK (light) |
| AMBP | 839.881424 | 2 | C[+57.021464]VLFPYGGC[+57.021464]QGNGNK (heavy) |
| ST6GAL1 | 397.213777 | 2 | FSAEALR (light) |
| ST6GAL1 | 402.217912 | 2 | FSAEALR (heavy) |
| ST6GAL1 | 539.781698 | 2 | C[+57.0]AVVSSAGSLK (light) |
| ST6GAL1 | 543.788798 | 2 | C[+57.0]AVVSSAGSLK (heavy) |
| FN1 | 742.35355 | 2 | GEWTC[+57.0]IAYSQLR (light) |
| FN1 | 747.357685 | 2 | GEWTC[+57.0]IAYSQLR (heavy) |
| FN1 | 955.725328 | 3 | C[+57.0]DPVDQC[+57.0]QDSETGTFYQIGDSWEK (light) |
| FN1 | 958.396727 | 3 | C[+57.0]DPVDQC[+57.0]QDSETGTFYQIGDSWEK (heavy) |
| FN1 | 585.228776 | 2 | YQC[+57.0]YC[+57.0]YGR (light) |
| FN1 | 590.23291 | 2 | YQC[+57.0]YC[+57.0]YGR (heavy) |
| BMP10 | 446.766177 | 2 | LYTLVQR (light) |
| BMP10 | 451.770312 | 2 | LYTLVQR (heavy) |
| BMP10 | 589.836996 | 2 | ITIFEVLESK (light) |
| BMP10 | 593.844096 | 2 | ITIFEVLESK (heavy) |
| HGF | 776.37172 | 3 | C[+57.0]EGDTTPTIVNLDHPVISC[+57.0]AK (light) |
| HGF | 779.04312 | 3 | C[+57.0]EGDTTPTIVNLDHPVISC[+57.0]AK (heavy) |
| HGF | 481.258716 | 2 | ESWVLTAR (light) |
| HGF | 486.262851 | 2 | ESWVLTAR (heavy) |
| THBS4 | 819.412796 | 3 | GAGSLELYLDC[+57.0]IQVDSVHNLPR (light) |
| THBS4 | 822.748885 | 3 | GAGSLELYLDC[+57.0]IQVDSVHNLPR (heavy) |
| THBS4 | 548.6299 | 3 | AFAGPSQKPETIELR (light) |
| THBS4 | 554.637389 | 3 | AFAGPSQKPETIELR (heavy) |
| THBS4 | 623.837528 | 2 | KPQDFLEELK (light) |
| THBS4 | 631.851727 | 2 | KPQDFLEELK (heavy) |
| THBS4 | 1086.522573 | 3 | GSLFQVASLQDC[+57.0]FLQQSEPLAATGTGDFNR (light) |
| THBS4 | 1089.858663 | 3 | GSLFQVASLQDC[+57.0]FLQQSEPLAATGTGDFNR (heavy) |
| THBS4 | 572.001445 | 3 | KPQDFLEELKLVVR (light) |
| THBS4 | 580.680334 | 3 | KPQDFLEELKLVVR (heavy) |
| JAG2 | 589.965065 | 3 | DHVPQGTTVGAIC[+57.0]SGIR (light) |

| | | | |
|---|---|---|---|
| JAG2 | 593.301155 | 3 | DHVPQGTTVGAIC[+57.0]SGIR (heavy) |
| JAG2 | 501.294245 | 2 | LLVLLC[+57.0]DR (light) |
| JAG2 | 506.298379 | 2 | LLVLLC[+57.0]DR (heavy) |
| JAG2 | 846.931208 | 2 | ASSGASAVEVAVSFSPAR (light) |
| JAG2 | 851.935342 | 2 | ASSGASAVEVAVSFSPAR (heavy) |
| PLG | 897.413479 | 2 | C[+57.0]TTPPPSSGPTYQC[+57.0]LK (light) |
| PLG | 901.420579 | 2 | C[+57.0]TTPPPSSGPTYQC[+57.0]LK (heavy) |

Cox, J., and Mann, M. (2008). MaxQuant enables high peptide identification rates, individualized p.p.b.-range mass accuracies and proteome-wide protein quantification. Nat. Biotechnol. *26*, 1367–1372.
Edfors, F., Boström, T., Forsström, B., Zeiler, M., Johansson, H., Lundberg, E., Hober, S., Lehtiö, J., Mann, M., and Uhlen, M. (2014). Immunoproteomics Using Polyclonal Antibodies and Stable Isotope–labeled Affinity-purified Recombinant Proteins. Mol. Cell. Proteomics *13*, 1611–1624.
Edfors, F., Forsstrom, B., Vunk, H., Kotol, D., Fredolini, C., Maddalo, G., Svensson, A.-S., Bostrom, T., Tegel, H., Nilsson, P., et al. (2018). Screening a Resource of Recombinant Protein Fragments for Targeted Proteomics. BioRxiv 472662.
Maclean, B., Tomazela, D.M., Shulman, N., Chambers, M., Finney, G.L., Frewen, B., Kern, R., Tabb, D.L., Liebler, D.C., and Maccoss, M.J. (2010). Skyline : an open source document editor for creating and analyzing targeted proteomics experiments. *26*, 966–968.
Tegel, H., Steen, J., Konrad, A., Nikdin, H., Pettersson, K., Stenvall, M., Tourle, S., Wrethagen, U., Xu, L.L., Yderland, L., et al. (2009). High-throughput protein production - Lessons from scaling up from 10 to 288 recombinant proteins per week. Biotechnol. J. *4*, 51–57.
Zeiler, M., Straube, W.L., Lundberg, E., Uhlen, M., and Mann, M. (2012). A Protein Epitope Signature Tag (PrEST) library allows SILAC-based absolute quantification and multiplexed determination of protein copy numbers in cell lines. Mol. Cell. Proteomics *11*, 1–13.
